# Long-term imaging of the ventral nerve cord in behaving adult *Drosophila*

**DOI:** 10.1101/2021.10.15.463778

**Authors:** Laura Hermans, Murat Kaynak, Jonas Braun, Victor Lobato Ríos, Chin-Lin Chen, Semih Günel, Florian Aymanns, Mahmut Selman Sakar, Pavan Ramdya

**Author notes:** equal contribution (Laura Hermans, Murat Kaynak).

## Abstract

The dynamics and connectivity of neural circuits continuously change during an animal’s lifetime on timescales ranging from milliseconds to days. Therefore, to investigate how biological networks accomplish remarkable cognitive and behavioral tasks, minimally invasive methods are needed to perform repeated measurements, or perturbations of neural circuits in behaving animals across time. Such tools have been developed to investigate the brain but similar approaches are lacking for comprehensively and repeatedly recording motor circuits in behaving animals. Here we describe a suite of microfabricated technologies that enable long-term, minimally invasive optical recordings of the adult *Drosophila melanogaster* ventral nerve cord (VNC)—neural tissues that are functionally equivalent to the vertebrate spinal cord. These tools consist of (i) a manipulator arm that permits the insertion of (ii) a compliant implant into the thorax to expose the imaging region of interest; (iii) a numbered, transparent polymer window that encloses and provides optical access to the inside of the thorax, and (iv) a hinged remounting stage that allows gentle and repeated tethering of an implanted animal for two-photon imaging. We validate and illustrate the utility of our toolkit in several ways. First, we show that the thoracic implant and window have minimal impact on animal behavior and survival while also enabling neural recordings from individual animals across at least one month. Second, we follow the degradation of chordotonal organ mechanosensory nerve terminals in the VNC over weeks after leg amputation. Third, because our tools allow recordings of the VNC with the gut intact, we discover waves of neural population activity following ingestion of a high-concentration caffeine solution. In summary, our microfabricated toolkit makes it possible to longitudinally monitor anatomical and functional changes in premotor and motor neural circuits, and more generally opens up the long-term investigation of thoracic tissues.

## 1 Introduction

Neural tissues are remarkably plastic, adapting to changes in internal states (e.g., injury, or hunger) and repeated exposure to salient environmental cues (e.g., during learning). In neuroscience, physiological studies of long timescale phenomena (e.g., memory formation and neurodegeneration) have often compared data pooled across animals sampled at different time points. However, resolving differences across conditions using this approach suffers from inter-individual variability. Thus, longitudinal recordings of the same animal are ideal for uncovering changes in neural dynamics and circuit reorganization. Important technical challenges must be overcome to perform long-term investigations of individual animals, including minimizing experimental insults and making them more tolerable.

With the advent of microscopy-based neural recordings, most notably two-photon calcium imaging [1], it has become possible to chronically record brain circuits *in vivo* in a minimally invasive manner. Cranial window technologies were first developed to study mouse neocortex [2] and have since been improved to acquire larger [3] and deeper [4] imaging fields-of-view, and longer duration recordings [5]. Similarly, long-term neural imaging has emerged as a promising tool for studying brain dynamics in the adult fly *Drosophila melanogaster* [6]. *Drosophila* offer the advantages of (i) being genetically tractable, (ii) having a small nervous system with many fewer neurons, and (iii) nevertheless generating complex social, navigation, and motor behaviors [7–10]. Building upon methods for recording brain circuits during behavior [11, 12], recent approaches also enable chronic recording of the fly brain [13, 14].

However, these techniques have been restricted to the study of superficial brain regions. Only very recently, has it become possible to image the activity of premotor and downstream motor circuits in the spinal cord of tethered, behaving mice [15, 16], and in the ventral nerve cord (VNC) of tethered, behaving flies [17]. The VNC is coarsely organized like the mammalian spinal cord [18], and its control principles also resemble those used by vertebrates—including the roles of central pattern generators (CPGs) and limb mechanosensory feedback [19, 20]. These features of the *Drosophila* VNC, including its relatively small size and genetic accessibility, make it an exceptionally promising model for the comprehensive investigation of motor circuit function across long time scales.

The VNC rests ventrally within the fly’s thorax beneath layers of opaque tissue including—from ventral to dorsal—salivary glands, gut, fat bodies, indirect flight muscles, and cuticle. Thus, until recently, it has not been possible to record from this neural tissue in tethered, behaving animals. We developed an approach that affords optical access to the VNC during behavior by surgically (and genetically, in the case of indirect flight muscles) removing these tissues [17]. However, this operation is invasive, requiring the resection of thoracic organs and leaving open the thoracic cavity. These interventions preclude recordings that last beyond a few hours and make repeated measurements of the same animal impossible. Thus, although this technique permits acute neural recordings of *Drosophila* premotor and motor circuits, understanding how these circuits can reorganize and adapt across time has remained out of reach.

Here, we describe a suite of microfabricated tools that permit long-term and repeated recordings of the *Drosophila* VNC for more than one month. These tools were inspired by those used to perform experiments on larger animals (e.g., cranial windows [3, 5] and implantable microprisms [4]) but were radically modified to address the unique challenges associated with studying extremely small animals (the fly is *∼*2-3 mm long). We used microfabrication approaches to construct tools that are orders of magnitude smaller and that permit extremely gentle tissue manipulations. Specifically, we designed (i) a manipulator (‘arm’) that allows us to move aside and temporarily hold in place thoracic organs, (ii) flexible, implantable structures that eliminate the need to surgically remove thoracic organs to access the VNC, (iii) a transparent polymer window that encloses the thoracic cavity and is numbered, allowing individual flies to be distinguished from one another across imaging sessions, and (iv) a remounting stage that allows flies to be gently yet firmly tethered, to perform repeated imaging of the same animal. We provide detailed descriptions of how to fabricate and use all of these tools, with the aim of enabling their adoption by other laboratories.

We illustrate the capabilities of this long-term imaging toolkit through a series of proof-of-concept studies. First, we demonstrate that implants and windows have minimal impact on animal survival and locomotor behavior, and that they permit neural recordings for at least one month. Second, we follow the degradation of limb mechanosensory neuron innervation of the VNC over two weeks after leg removal. Third, we illustrate how—by leaving thoracic organs intact—one can investigate the influence of drug ingestion on neural dynamics. Taken together, these studies illustrate how our long-term thoracic imaging toolkit enables the discovery of changes in neural morphology and activity over time. These tools may be applied to perform longitudinal investigations of other thoracic tissues as well including indirect flight muscle, fat bodies, gut, and trachea.

## 2 Results

### 2.1 Long-term recording toolkit and experimental workflow

We developed microfabricated technologies and a micromanipulation protocol that enable optical access to the fly’s VNC for more than one month. Implanted flies exhibit no obvious deficits in their ability to feed, walk, lay eggs, or interact with others. **(Figure 1A)(Video 1)**. This toolkit consists of two major components: a compliant and transparent implant **(Figure 1B)** and a numbered, transparent thoracic window **(Figure 1C)**. The implants are fabricated *en masse* using soft lithography, a technique that is based on rapid prototyping and replica molding **(Figure S2** and **Figure S3)**. The window is fabricated from a biocompatible polymer, SU-8, using conventional photolithography **(Figure S1)**. To use these tools, we developed a manipulation protocol illustrated in **Video 2**. Briefly, we first mount animals onto a surgical dissection stage using UV-curable glue **(Figure 1D)** [17]. Next, we cut a square-shaped hole into the dorsal cuticle using a 30G syringe needle **(Figure 1E-i)**. The indirect flight muscles (IFMs) were subsequently removed to create a thoracic opening for the implant. To minimize the impact of the microsurgery, we worked with animals expressing the apoptosis-inducing protein, Reaper, specifically in IFMs (*Act88F:Rpr*). Expressing Reaper results in rapid degradation of the muscle tissue [17], the remainder of which can easily be removed with the syringe needle. Having exposed the thoracic tissues, we used a fine glass needle and forceps to unilaterally detach tracheal fibers that connect the gut and left salivary gland. We designed a custom manipulation arm **(Figure S4)** to push the internal organs—gut, salivary gland and trachea—to the right side of the thoracic cavity **(Figure 1E-ii)** and insert the implant, in a closed state, into the available space **(Figure 1E-iii)(Figure S3)**. Upon release, the implant gradually opened, holding the organs against the thoracic wall after the manipulation arm was retracted **(Figure 1E-iv)**. We sealed the exposed thoracic cavity by gluing a transparent polymer window to the cuticle **(Figure 1E-v)**. These windows have unique numbers engraved on their surfaces, making it possible to identify and distinguish between implanted animals. By removing the UV-curable glue holding the animal’s scutellum to the dissection stage, we could then detach animals, allowing them to behave freely.

**Figure 1:**
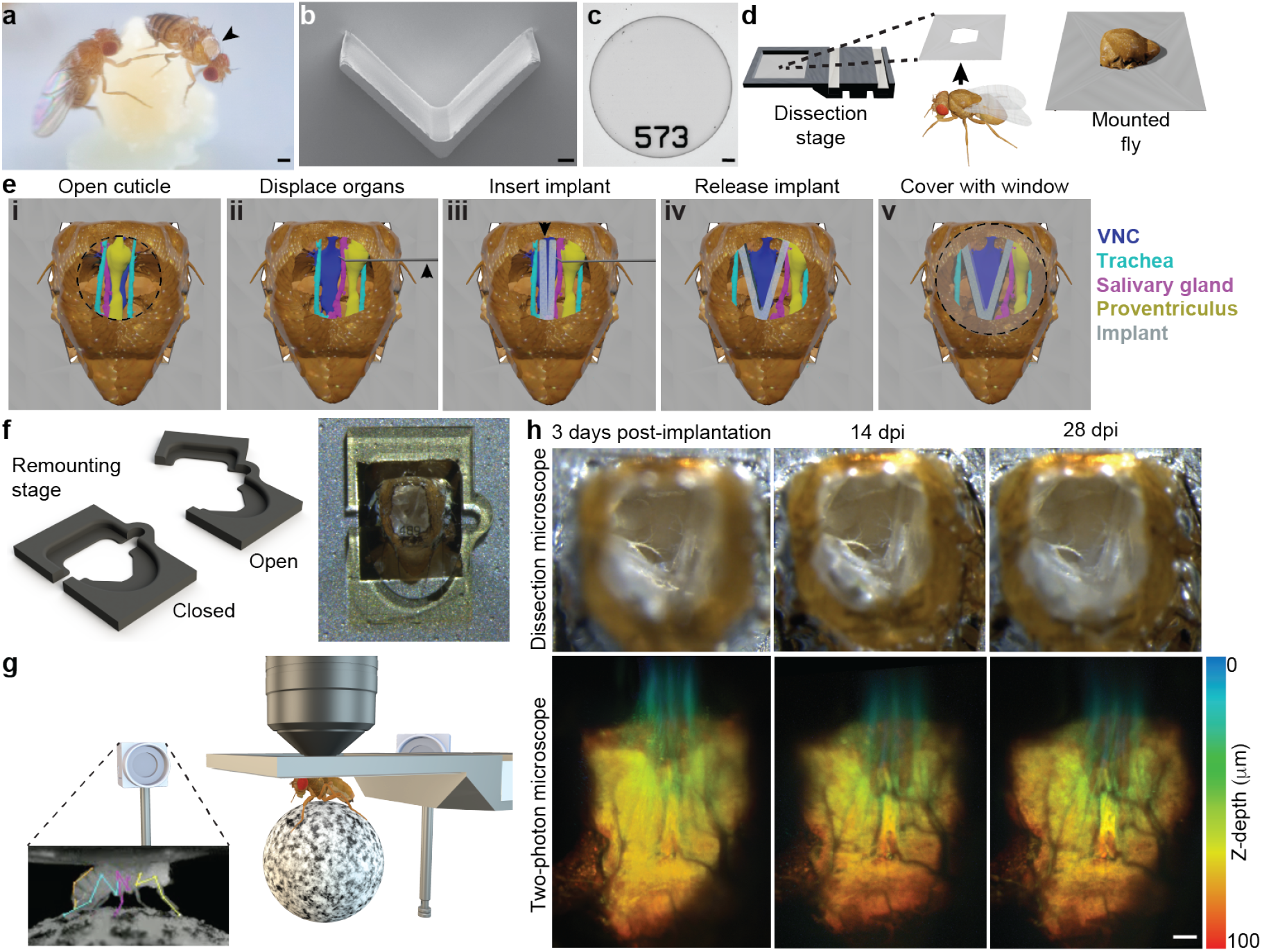
Long-term recording technologies, workflow, and experimental validation. **(A)** Implanted adult flies can be raised in complex environments between neural recordings. Here an implanted animal—see dorsal thoracic window (black arrow)—interacts with a non-implanted animal. Scale bar is 0.5mm. **(B)** A mechanically compliant and transparent implant microfabricated from Ostemer 220. Scale bar is 50 μm. **(C)** A numbered, transparent thoracic window microfabricated from SU-8. Scale bar is 50 μm. **(D)** For implantation, an animal is first mounted, thorax first, into a hole in a steel shim within a dissection stage. **(E)** A multi-step dissection permits long-term optical access to the ventral nerve cord (VNC). **(i)** First, a hole is cut into the dorsal thoracic cuticle, revealing the proventriculus (yellow), trachea (cyan), and salivary gland (magenta) overlying the ventral nerve cord (VNC, dark blue). The indirect flight muscles (IFMs) were degraded by tissue-specific expression of Reaper (*Act88F:Rpr*) [17]. **(ii)** Then, using a custom-designed manipulator arm, thoracic organs are displaced, revealing the VNC. **(iii)** Next, the implant is placed within this thoracic hole in a narrow, mechanically closed configuration. **(iv)** The arm is then removed and the implant is released, causing it to open and mechanically push aside organs covering the VNC. **(iv)** Finally, a transparent window is sealed to enclose the thoracic hole. **(F)** A remounting stage permits gentle mounting and dismounting of animals for repeated two-photon imaging. **(left)** A flexible microfabricated hinge allows the stage to open and close. **(right)** Sample image of an animal tethered to the remounting stage as seen from above. **(G)** Implanted animals tethered to the remounting stage are placed under a two-photon microscope surrounded by a camera array. This configuration permits simultaneous recordings of neural activity and animal behavior. Inset shows one camera image superimposed by deep learning-based 2D poses estimated using DeepFly3D [21]. **(H) (top row)** The dorsal thorax of an implanted animal, as seen from the dissection microscope, and **(bottom row)** its VNC, as visualized using the two-photon microscope. This animal expresses GFP throughout the nervous system and is recorded at **(left)** 3 dpi, **(middle)** 14 dpi, and **(right)** 28 dpi. Z-stacks are depth color-coded (100 μm). Scale bar is 25 μm.

To facilitate repeated neural imaging of implanted flies, we printed a remounting stage **(Figure 1F)(Figure S5)** using two-photon polymerization. This manufacturing process has the accuracy required to fabricate 3D features that reliably hold animals in place. When mounted, animals were studied using a two-photon microscope surrounded by a multi-camera array. This system enables simultaneous recordings of neural activity in the VNC [17] as well as markerless 3D body part tracking [21] **(Figure 1G)**. In the vast majority of cases, our implantation protocol was successful. Infrequently, implanted animals exhibited specific movements of respiratory, or digestive tissues that could occlude the VNC during imaging **(Figure S6)**. Successful implantation permitted optical access to the VNC that remained largely unchanged over one month and allowed repeated studies of the structure **(Video 3) (Figure 1H)** and functional dynamics of neural circuits **(Video 4)**.

### 2.2 Impact of long-term imaging technologies on lifespan and behavior

Next, we studied the potential impact of implantation on animal lifespan. Specifically, we measured the longevity of three groups of animals (n = 40 per group): flies that (i) were not manipulated (‘Intact’), (ii) endured cold anesthesia, mounting onto the dissection stage, and wing removal (‘Sham dissected’), or (iii) underwent the full implantation procedure (‘Implanted’). For this experiment, 73% of implanted animals survived surgery. We observed that implanted flies could survive up to 88 days, but had a more rapid mortality rate than intact animals in the first days following implantation **(Figure 2A)**. Notably, sham implanted flies also had increased mortality in those first days, suggesting that pre-implantation animal handling, and not the implantation procedure, was responsible.

**Figure 2:**
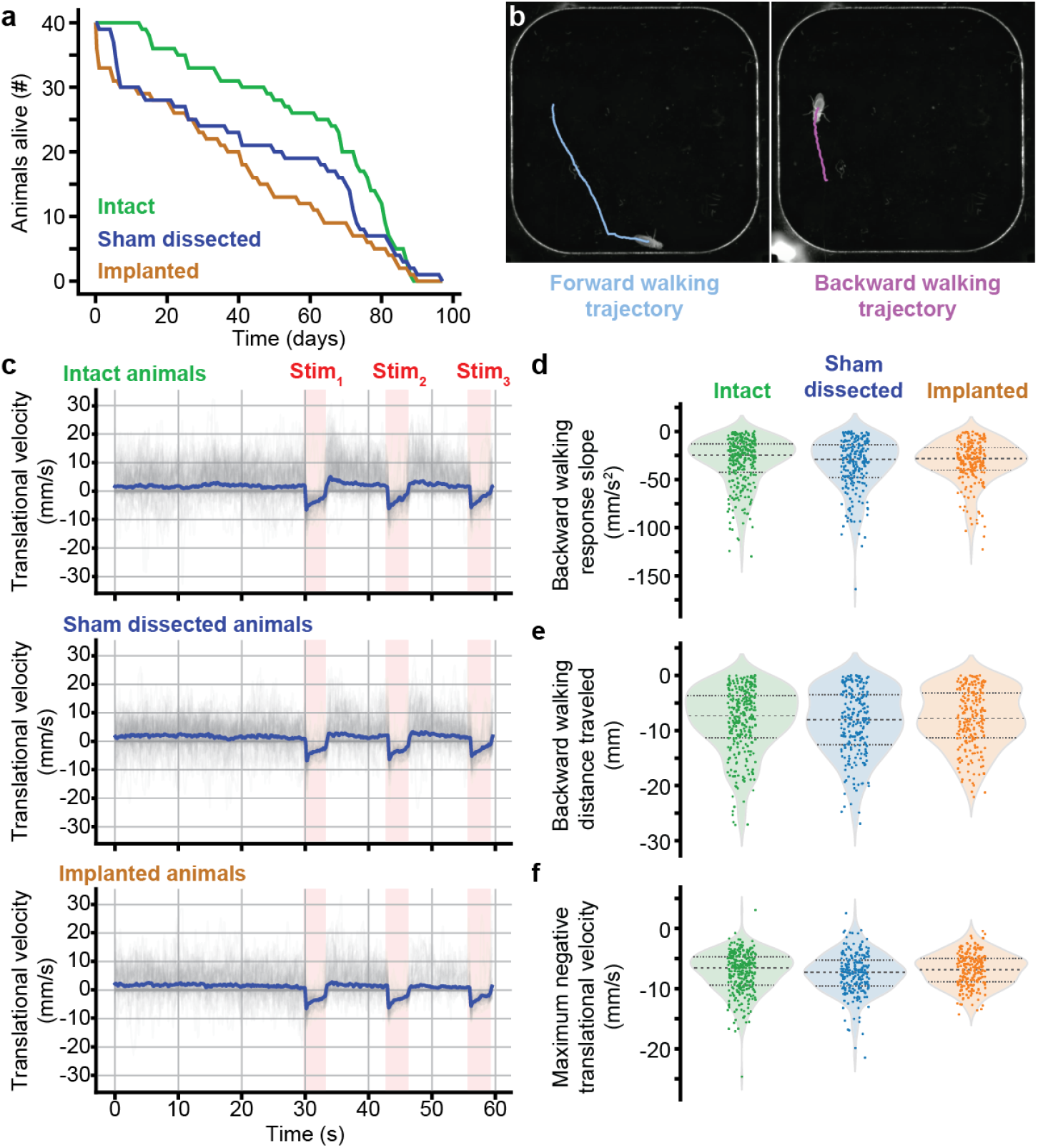
Impact of long-term imaging technologies on lifespan and behavior. **(A)** Survival curves for genetically-identical sibling animals that were (i) not experimentally manipulated (green, ‘Intact’), (ii) tethered, cold anaesthetised, and had their wings removed (blue, ‘Sham dissected’), or (iii) prepared for long-term imaging by implantation and the addition of a thoracic window (orange, ‘Implanted’). **(B)** Behaviors were compared by analyzing the dynamics of optogenetically activated backward walking within a rounded square-shaped arena. Locomotion was computationally analyzed and plotted, showing the animal’s initial forward trajectory (cyan) and subsequent optically evoked backward walking trajectory (purple). **(C)** Translational velocities of intact **(top)**, sham dissected **(middle)**, and implanted **(bottom)** animals during 30 s of spontaneous behavior, followed by three optogenetic stimulation periods of 3 s each (pink, ‘Stim’). Shown are the raw (grey) and mean (blue) traces. From these time-series, we calculated summary statistics including **(D)** the initial negative slope in translational velocity—backward walking—upon optogenetic stimulation, **(E)** the integrated translational velocity over the entire optogenetic stimulation period, and **(F)** the peak negative translational velocity over the entire optogenetic stimulation period.

Although implantation did not dramatically affect longevity, placing a microfabricated object within the thorax might negatively impact walking, possibly due to the perturbation of leg-related musculature, or simply the additional weight. Investigating this possibility is difficult because of the variety of gaits used at different walking speeds and maneuvers. Therefore, to be able to perform quantitative analysis of kinematics, we investigated the stereotyped backward walking response of flies to optogenetic activation of Moonwalker Descending Neurons (MDNs) [22]. Specifically, we stimulated animals expressing the light-gated cation channel, CsChrimson [23], in MDNs [24] repeatedly over the course of one month. We analyzed the trajectories of intact, sham implanted, or implanted flies walking in a custom-built arena **(Figure 2B)**. We first recorded spontaneous behaviors for 30 s, and then delivered three consecutive flashes of orange light for 3 s each **(Figure 2C, pink)** with an inter-stimulus interval of 10 s. Upon optogenetic stimulation, animals generated fast backward walking that gradually slowed and rapidly returned to baseline when the light was turned off **(Video 5)**. Over all recording sessions, we did not measure any significant difference in the translational velocities for intact, sham dissected, and implanted animals in terms of the initial backward acceleration **(Figure 2D)** (*P* =0.31; Kruskal-Wallis test), the total backward walking distance traveled **(Figure 2E)** (*P* =0.80; Kruskal-Wallis test), and the maximum backward walking velocity **(Figure 2F)** (*P* =0.27; Kruskal-Wallis test). Similar results were obtained when comparing age-restricted cohorts, aside from a small difference in maximum negative translational velocity for sham dissected animals at 14-16 dpi compared with the other two groups **(Figure S7)**. Taken together, these results suggest that locomotion is not significantly impacted by the implantation procedure, and the presence of an additional thoracic payload.

### 2.3 Quantifying long-term structural degradation in the VNC following limb amputation

Neuronal circuits retain the capacity for structural rearrangement throughout adulthood [25,26]. This dynamism enables adaptive behavior even in the face of profound structural changes accompanying brain and spinal cord injury [27–29], or stroke [30]. Similarly, in flies, locomotor gaits reorganize following leg amputation [31] but the impact of this injury on locomotor circuits remains unexplored: uncovering associated changes in neural structures, or dynamics would require visualizing the VNC of amputated animals across days, or weeks. To illustrate how pur long-term imaging toolkit is ideally suited for these kinds of studies, we followed the degradation of primary proprioceptive mechanosensory afferents of an amputated leg. Specifically, we visualized the terminals of chordotonal organs (*Act88F-Rpr/+; iav-Gal4/UAS-GFP; +/+*) within the T1 (foreleg) VNC neuropil. Flies were implanted on the first day post-eclosion (dpe). Then, at one day post-implantation (1 dpi), we performed two-photon microscopy to acquire a 3D image volume of the VNC, consisting of 100 images at 1 μm depth intervals. Then, at 2 dpi, the front left leg of each experimental animal was amputated near the thorax-coxa joint **(Figure 3A)**.

**Figure 3:**
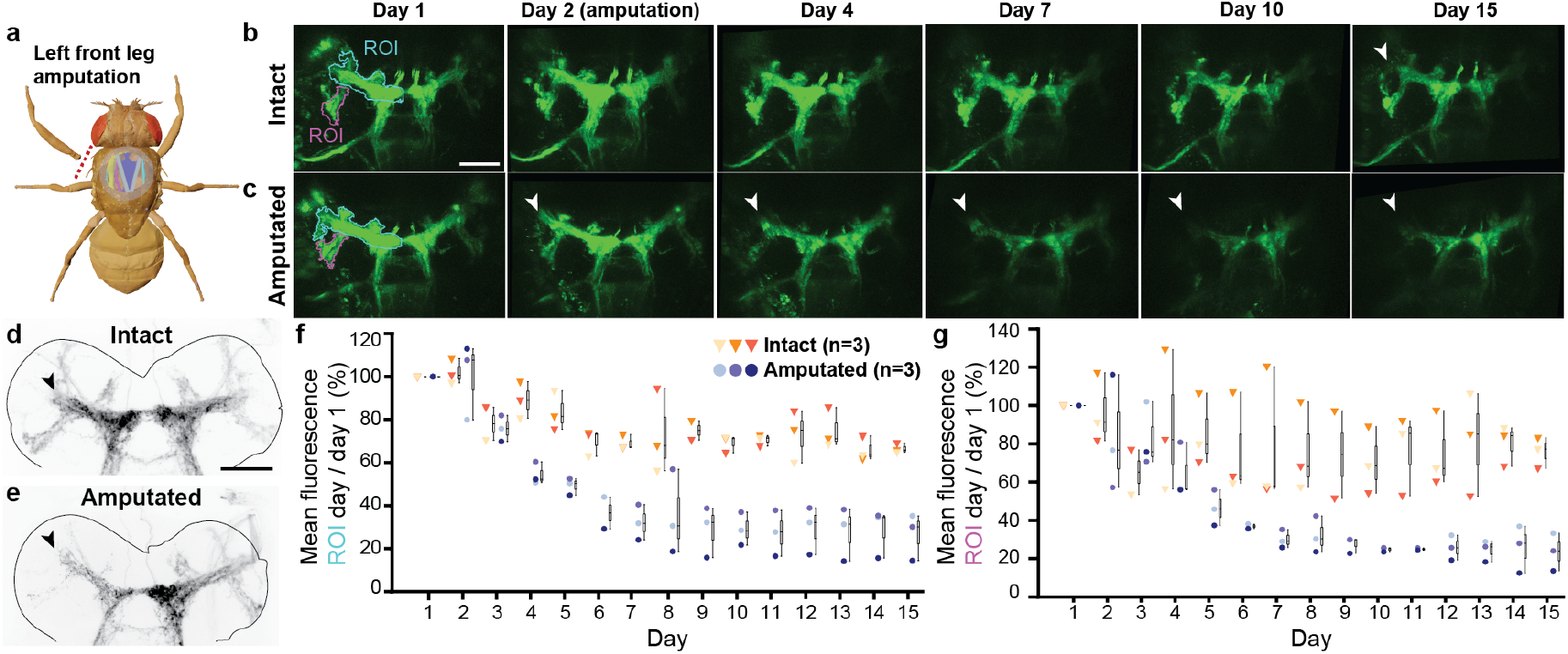
Long-term imaging of mechanosensory nerve degradation in the VNC following limb amputation. **(A)** In experimental animals, the front left leg was amputated at the thorax-coxa joint at 2 dpi. **(B,C)** Maximum intensity projections of z-stacks recorded from **(B)** an intact (control), or **(C)** front left leg amputated animal. Data were acquired using two-photon microscopy of an implanted animal. Shown are images taken at 1, 2, 4, 7, 10 and 15 dpi. Images are registered to the 1 dpi image. Scale bar is 50 μm. White arrowheads indicate degrading axon terminals in the VNC. **(D,E)** Standard deviation projections of confocal z-stacks recorded from dissected and stained VNCs (nc82 staining—not shown—is outlined in grey, GFP fluorescence is black). Tissues were taken from implanted animals whose front left legs were either **(D)** left intact, or **(E)** amputated at 2 dpi. VNC tissue was removed and stained at 20 dpi. Black arrowhead indicates VNC region exhibiting greatest difference between intact and amputated proprioceptor innervation. Scale bar is 50 μm. **(F,G)** Fluorescence measured across days using two-photon microscopy from intact animals (n=3; orange-red triangles), or animals whose front-left legs were amputated at 2 dpi (n=3; blue circles). Measurements indicate mean fluorescence within the **(F)** blue, or **(G)** pink region-of-interest (ROI) as in panels B and C, normalized and divided by the mean fluorescence at 1 dpi. Box plots indicate median, upper, and lower quartiles.

Implanted flies tolerated leg amputation, and displayed normal behavior with five legs (data not shown). Every day for 15 days, we collected image volumes of the VNC’s T1 neuropil from control flies (‘Intact’) and those with their front left leg removed (‘Amputed’). In control animals, we observed some photobleaching throughout the imaging region over days **(Figure 3B)**. However, the decline in fluorescence intensity was not nearly as profound as the signal reduction observed among chordotonal organ axon terminals in the left T1 neuropil of leg amputated animals **(Figure 3C)(Video 6)**. Posthoc confocal imaging confirmed that T1 left leg sensory innervation of the VNC persisted in intact animals **(Figure 3D)** but degraded in amputated animals **(Figure 3E)**. By quantifying changes in signal intensity within specific regions of interest (ROIs) of chordotonal axon innervations of the VNC [32], we observed a marked reduction in the fluorescence over time that was highly reproducible **(Figure 3F,G) (Figure S8)**.

### 2.4 Capturing neural population dynamics associated with caffeine ingestion

In addition to being morphologically adaptable across days and weeks, neural circuits also continuously modulate their dynamics on shorter timescales (e.g., minutes to hours) depending on the internal state of the animal. In *Drosophila*, as in vertebrates, these states naturally vary with hunger [33], sleep [34], sexual arousal [35], aggression [36], and defensive arousal [37]. Internal states can also change following the ingestion of psychoactive substances like caffeine [38–40]. Continuous monitoring of the nervous system will be instrumental to understand how neural circuits reconfigure when animals switch from one state to another.

The previous technique for studying VNC neural dynamics in behaving animals [17] required the removal of large sections of gut, reducing the longevity of animals and, thus, making hours or days-long experiments that study hunger and sleep states impossible. Furthermore, removing the gut precludes feeding, and, consequently, does not allow one to investigate how satiety, or ingesting psychoactive substances influences neural dynamics. Here we aimed to demonstrate how our long-term imaging technology, which preserves the gut, allows animals to be fed during two-photon microscopy, and, therefore, enables the interrogation of how drug intake modulates neural dynamics. Specifically, we explored the impact of high concentrations of caffeine on global brain states, as measured by the activity of ascending and descending neuron populations whose axons pass through the thoracic cervical connective. To do this, we recorded a coronal cross-section of the connective [17] in flies expressing the genetically encoded calcium indicator, GCaMP6f, as well as the anatomical marker, tdTomato, throughout the nervous system (*Act88F-Rpr/+; GMR57C10-Gal4/UAS-opGCaMP6f; UAS-tdTomato/+*). We could resolve the activity levels of hundreds of neurons including higher-order integrative descending neurons that drive actions [41, 42], and ascending neurons that convey behavioral state to the brain [43].

Previous studies have shown that flies exposed to low doses of caffeine have reduced sleep [38, 39] and increased locomotor activity [40]. We asked to what extent caffeine ingestion would change global neural dynamics. We starved animals for 21-23 h to encourage feeding. Then, after implantation, we recorded neural activity in the cervical connective (‘Before feeding’). While continuing to image, animals were then fed **(Figure 4A)** either a control solution (‘Sucrose only’) containing 8 mg/ml sucrose and 1 mg amaranth dye (to confirm feeding [44]) **(Video 7)**, or an experimental solution that also contained 8 mg/ml, or 40 mg/ml caffeine: ‘Low caffeine’ **(Video 8)**, or ‘High caffeine’ **(Video 9)**, respectively. We continued to record neural activity and behavior for the next 32 minutes. Feeding was confirmed by posthoc evaluation of abdominal coloration due to dye ingestion **(Figure 4B)**.

**Figure 4:**
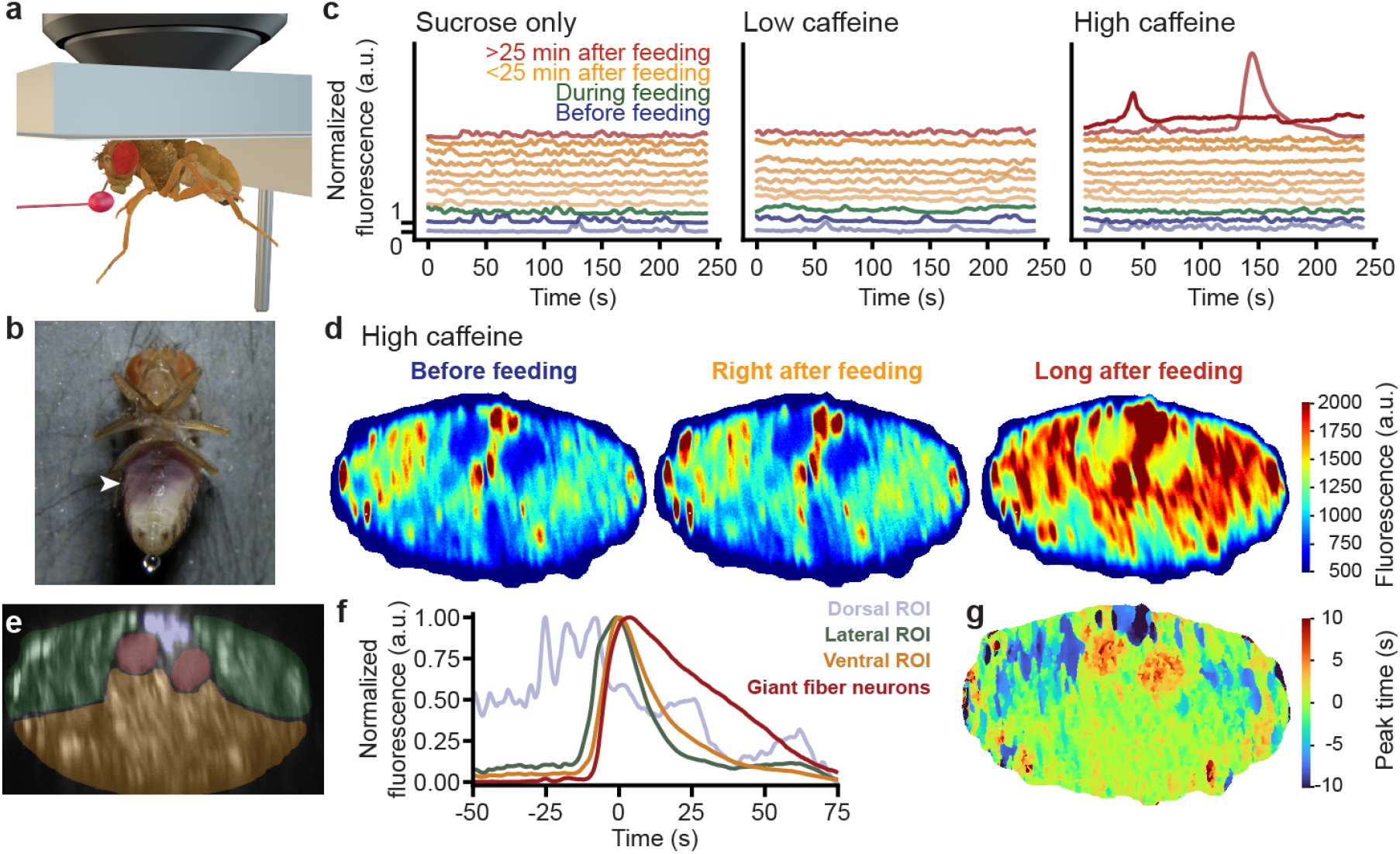
Continuous imaging of neural population dynamics before, during, and after caffeine ingestion. **(A)** Digital rendering of a fly being fed while neurons are recorded using a two-photon microscope. **(B)** Photo of an implanted animal after ingesting a high-concentration caffeine solution during two-photon imaging. White arrowhead indicates purple coloration of the abdomen, confirming digestion of a caffeine-sucrose solution mixed with Amaranth dye. **(C)** Normalized fluorescence across all axons passing through the thoracic neck connective during four minute recordings either before (blue), during (green), soon after (orange), or long after (red) feeding. Flies were fed with a solution containing either only sucrose (left), sucrose and a low-dose (middle), or high-dose of caffeine (right). **(D)** Color-coded mean neural activity during all non-locomotor periods for a fly either before (left), immediately after (middle), or long after (right) ingestion of a high-concentration caffeine solution. **(E)** The cervical connective in one implanted animal is segmented into four regions-of-interest (ROIs). These are overlaid on a standard-deviation time-projection image. **(F)** Neural activity normalized to peak fluorescence during a wave of activity. Traces are color-coded as in panel E. The peak of mean fluorescence across all regions is centered on 0 s. **(G)** Pixel-wise time of peak activity. The peak of mean activity across the entire neck connective set as 0 s.

Across all three experimental conditions—before, during, and shortly after feeding—we observed fluctuations in neural activity that were largely associated with epochs of walking and grooming **(Figure 4C, blue, green, and orange traces; Videos 7-9)**. However, more than 25 min after feeding, we observed large waves of activity in the high caffeine condition **(Figure 4C, red traces)**. Waves were much larger in amplitude (up to 800%Δ*F/F*) than activity associated with behaviors like walking and grooming (up to 200%Δ*F/F*). The wave spread across the connective **(Figure 4D)** and was associated with a rigid pose accompanied by micromovements **(Video 10)**. We could observe these waves several times in animals fed a high caffeine solution and they were observed in all animals **(Figure S9)**. The temporal evolution of caffeine-induced waves were also reproducible **(Figure 4E)**. Neurons were active in the spatial order: dorsalmedial (blue), dorsolateral (green), and then ventral (orange) connective. Finally, the giant fibers (red) [45] became active and sustained this activity over longer timescales **(Figure 4F,G)**. These data illustrate how, in addition to long-term studies of sleep or learning, our long-term imaging technology enables the investigation of how food or drug intake influences internal states and global neural dynamics.

## 3 Discussion

Here we have described a microfabricated toolkit that enables long-term imaging of tissues in the adult *Drosophila* thorax including premotor and motor circuits in the VNC. Our toolkit consists of (i) a micromanipulator arm, (ii) a polymer-based soft implant that displaces thoracic organs, (iii) a numbered, transparent polymer window that seals the thoracic opening, and (iv) a compliant tethering stage that permits repeated mounting of animals for two-photon imaging. Taken together, these tools expand the neural recording window from a few hours [17] to more than one month without markedly reducing the lifespan of implanted animals, or significantly perturbing their locomotor behavior. We illustrated several use cases for our long-term imaging approach including (i) recording neural morphology **(Video 3)** and function **(Video 4)**, (ii) recording the weeks-long degradation of proprioceptive neurons from an amputated limb, and (iii) uncovering global waves of neural activity following caffeine ingestion.

Our longevity experiment showed that the total lifespan of implanted flies was similar to that of intact flies. The survival curves were, however, shifted for implanted and sham dissected flies due to an excess mortality within the first few days following surgery. This suggests that those initial losses might be due to surgical handling and not specifically linked to implantation. Consistent with this, our studies of backward walking demonstrated no clear changes in several locomotor metrics. However, in the future, it would be worth analyzing the impact of implants on more complex behaviors like courtship and copulation.

While recording the anatomy of chordotonal projections to the VNC over two weeks, we observed a marked reduction in fluorescence in the T1 neuromere in the first week following leg amputation. Although relatively stable for some regions-of-interest, this reduction continued for others. This heterogeneity is consistent with the fact that these terminals arrive from distinct chordotonal cell populations [32] which may have varying levels of robustness against degradation. Alternatively, some terminals might also arise via ascending projections from T2 (midleg) or T3 (hindleg) and thus not be affected by foreleg amputation.

While visualizing the activity of descending and ascending neurons in the thoracic cervical connective after caffeine ingestion, we did not observe large changes in neural activity after ingestion of the low-concentration caffeine solution, despite reported behavioral changes. [40] On the other hand, we discovered large waves of activity following ingestion of a high-concentration caffeine solution. Some flies exhibited several of these waves, suggesting that they are not due to calcium release during a terminal cell death process. However, the mechanistic basis and temporal propagation of these waves should be further investigated in future studies.

Based on these use case results, we envision that our microfabricated long-term imaging tools can be leveraged to study a variety of additional questions and challenges. For example, one might apply long-term VNC imaging to record the progression of neuronal loss in *Drosophila* models of disorders including Parkinson’s disease [46]. In their current form, our tools could also be used to enable drug screens of neural function in adult *Drosophila*. Additional steps might be taken to automate implantation by, for example, opening the thoracic cuticle using a UV-excimer laser [47], and developing robotic manipulation techniques to displace thoracic organs, position the implant, and seal the thoracic hole with a window in an automated fashion [48].

The implant fabrication pipeline is general; therefore, the form of the implant could be adapted to address other experimental challenges. For example, one might desire targeting specific tissues within the VNC such as circuits in the abdominal ganglia that regulate mating receptivity in females [49]. Furthermore, implants might be modified to store and release active components that, for example, deliver compounds into the hemolymph in a controlled manner.

In summary, our long-term imaging toolkit permits a variety of experiments on individual animals across a wide range of time scales, opening up the exploration of how biological systems—in particular premotor and motor circuits—adapt during aging or disease progression, following injury or learning, in response to changes in internal states and social experiences, and as a consequence of food or drug ingestion.

## 4 Materials and Methods

### 4.1 Fabrication of thoracic windows with engraved markers

Thoracic windows (transparent polymer disks) were fabricated using photolithography [50]. All exposure steps were performed on a mask aligner (MJB4, Süss MicroTec, Germany) using i-line illumination. Chrome masks were fabricated using a direct laser writer (VPG-200, Heidelberg Instruments, Germany) and an automatic mask processor (HMR900, HamaTech, Germany). The dimensions of microfabricated structures were measured using an optical microscope (DM8000 M, Leica Microsystems, Switzerland) or a mechanical surface profiler (Dektak XT, Bruker Corporation, USA). The protocol began with treating the surface of a 4-inch silicon wafer with a plasma stripper (PVA TePla 300, PVA AG, Germany) at 500 W for 5 min to reduce its wettability. An aqueous solutions of 25% (wt/vol) Poly(acrylic acid) (Polysciences, MW 50000) was spun at 2000 rpm (WS-650-23, Laurell Technologies Corporation, USA) to form a 1 μm thick sacrificial layer. This layer permits windows to be gently released at the end of the fabrication process **(Figure S1A-i)**. A negative photoresist (SU-8 3025, Kayaku Advanced Materials, USA) was directly spin-coated on the sacrificial layer and soft-baked **(Figure S1A-ii)**. After exposure, the windows were post-baked and uncured resist was removed with a developer (Propylene glycol methyl ether acetate (PGMEA, 1-methoxy-2-propanol acetate), Sigma-Aldrich, Germany) **(Figure S1A-iii)**. Next, the wafer with SU-8 windows was coated with a 20 μm thick layer of positive photoresist (AZ 40XT) using an automated processing system (ACS200 Gen3, Süss MicroTec, Germany). This extra layer of polymer serves as a physical mask during the metal deposition process. A second chrome mask was fabricated to pattern unique identifiers onto the windows using photolithography. Next, the wafer was coated with Ti and Au films [51] using physical vapor deposition (EVA 760, Alliance-Concept, France) at a thickness of 2 nm and 10 nm, respectively **(Figure S1A-iv)**. The development of the negative photoresist (Remover 1165, Kayaku Adv. Mat., USA) removed all the layers on top of the windows except for the numbers that serve as markers. Finally, the labelled windows were released by dissolving the sacrificial layer in DI water **(Figure S1A-vi)**. The windows were filtered, dried at room temperature, and sterilized prior to use in experiments. The resulting windows were optically transparent **(Figure S1B)** and of the appropriate size to seal thoracic openings **(Figure S1C)**.

### 4.2 Fabrication of polymer molds that are used to cast implants

We developed a two-level microfabrication technique to maximize throughput, protect master molds from excessive use, and facilitate technology dissemination [52, 53]. Briefly, implants were cast within elastomer templates that were fabricated from an etched wafer serving as a master mold. First, a four-inch silicon test wafer (100/P/SS/01-100, Siegert Wafer, Germany) was treated with hexamethyldisilazane (HMDS) (CAS number: 999-97-3, Sigma-Aldrich, Germany) and dehydrated at 125°C to enhance adhesion to its surface. The wafer was then spin-coated with an 8 μm thick film of positive photoresist (AZ 9260, Microchemicals GmbH, Germany) using an automatic resist processing system (EVG 150, EV Group, Germany)**(Figure S2A-i)**. After baking, exposure, and development steps, the wafer was then processed using deep reactive ion etching (DRIE), specifically a Bosch process, [54] (AMS 200 SE, Alcatel) to obtain nearly vertical walls with a high aspect ratio **(Figure S2A-ii)**. The remaining positive resist was stripped in a remover (Remover 1165, Kayaku Advanced Materials, USA) at 70°C and cleaned by rinsing with water and air drying **(Figure S2A-iii)**. The elastomer templates were fabricated by replica molding using polydimethylsiloxane (PDMS). The replica molding process began with vapor deposition of silane (trichloro(1H,1H,2H,2H-perfluorooctyl) Silane, Sigma-Aldrich, Germany) onto the surface of the master mold in a vacuum chamber for 6 h. Silanizion was only performed once because it forms a permanent silane layer. PDMS was prepared as a mixture (10:1, wt/wt) of the elastomer and the curing agent (GMID number: 01673921, Dow Europe GmbH, Germany) and poured onto the wafer in a petri dish. To release any bubbles trapped inside the high aspect ratio wells, the mold was degassed using a vacuum pump (EV-A01-7, Swiss Vacuum Technologies SA, Switzerland) in a vacuum desiccator (F42020-0000, SP Bel-Art Labware & Apparatus, USA). Finally, the elastomer was cured at 65°C for 5 h in an oven (UF30, Memmert GmbH, Germany) and the PDMS slab was peeled off **(Figure S2B)**. Using alignment markers as a guide, the slab was then cut into several pieces with a razor blade to serve as templates with which one could then fabricate implants **(Figure S2C)**.

### 4.3 Fabrication of implants

Flexible implants were fabricated from a photocurable polymer (Ostemer 220, Mercene Labs AB, Sweden). Polymerization occurs when a mixture of the base (Part B) and hardener (Part A) are exposed to UV light **(Figure S3A-i)**. The PDMS template was silanized (trichloro(1H,1H,2H,2H-perfluorooctyl) silane, Sigma-Aldrich, Germany) for 1 h in a vacuum desiccator **(Figure S3A-ii)**. Part A was warmed at 48°C overnight to make sure there were no undissolved crystals remaining in the solution. Part B and the container were also heated up to 48°C before mixing. Parts A and B were then mixed thoroughly and the mixture was degassed in a vacuum chamber for 5 min. A 200 μL drop of the mixture (1:1.86, wt/wt) was poured onto the template **(Figure S3A-iv)** and the template was mechanically sandwiched between two glass slides using two clips. The glass slide touching the implant polymer was previously plasma treated (PDC-32G, Harrick Plasma,USA) at 29 W for 1 min to facilitate implant release by improving the adhesion between the glass and implants. The solution was exposed to UV light (365 nm, UV9W-21, Lightning Enterprises, USA) for 10 min for polymerization **(Figure S3A-v)**. The samples were rotated several times during UV exposure to ensure a homogeneous reaction throughout the template. The implants were released by mechanically agitating the templates in isopropyl alcohol (IPA) using a sonicator (DT 100 H, Bandelin Sonorex Digitec, Germany)**(Figure S3A-vi)**. This whole process yielded a wafer with 100 implants **(Figure S3B,C)** that were subsequently cut out using a razor blade prior to implantation.

### 4.4 Fabrication of a manipulator arm that temporarily displaces thoracic organs

We designed and constructed a manipulator arm to temporarily displace thoracic organs during implantation **(Figure S4A,B)**. To construct the arm, we first 3D printed a mold that allowed us to glue a dissection pin (26002-10, Fine Science Tools, Germany) to the tip of a syringe needle (15391557, Fisher Scientific, USA) in a reproducible manner **(Figure S4C)**. The pin is inserted into the needle until its tip touches the end of the mold. We glued the pin to the needle using a UV-curable adhesive (Bondic, Aurora, ON Canada). The arm was then bent using forceps and guided by a second 3D printed mold **(Figure S4D)**. The pin was first bent coarsely and then adjusted more finely using the 3D printed mold. Another 3D printed piece was then used to connect the syringe needle to a 3-axis micromanipulator (DT12XYZ, ThorLabs, USA) and to an extension stage **(Figure S4A)**. The whole structure was then attached to a breadboard (MB1224, ThorLabs, USA) **(Figure S4B)**.

### 4.5 Fabrication of a remounting stage

We used direct laser writing [55] to fabricate a custom compliant mechanism that holds flies in place during two-photon microscopy. The mechanism was designed using 3D CAD software (SolidWorks 2021, Dassault Systèmes, France). A 25 mm x 25 mm diced silicon wafer was used as the substrate upon which structures were printed. The surface of the substrate was plasma treated at 500 W for 5 min and coated with an aqueous solution of 10% (wt/vol) Poly(acrylic acid) (MW 50000, Polysciences, USA) at 2000 rpm for 15 s using a spin-coater (WS-650-23, Laurell Technologies Corporation, USA) **(Figure S5A-i-iii)**. The mechanism was fabricated using a direct laser writer (Photonic Professional GT+, Nanoscribe GmbH, Germany) that controls two-photon polymerization **(Figure S5A-iv)**. A polymer (IP-S, Nanoscribe GmbH, Germany) was chosen as the print material due to its Young’s modulus of 4.6 GPa [56] and the resolution at which structures could be printed. The overall design was segmented into multiple frames because the maximum laser scan area provided by a 25X objective (NA 0.8, Zeiss) is 400 μm. This approach results in fine printing over a relatively large layout. The objective was dipped into liquid photoresist during printing. At the end of the printing process, the uncured polymer was removed using a developer (PGMEA, Sigma-Aldrich, Germany) for 20 min **(Figure S5A-v)**. Finally, the PGMEA was rinsed using IPA. The mechanism was released from the substrate by dissolving the sacrificial layer in DI water **(Figure S5A-vi)**. This yielded a microfabricated structure large enough to contain the thorax of the fly **(Figure S5B,C)**. The remounting stage was completed by attaching the mechanism onto a laser-cut aluminum frame using UV-curable glue (Bondic, Aurora, ON Canada).

### 4.6 Implantation procedure

The steps required to prepare flies for long-term VNC imaging are described here. See **Video 2** for more details.

#### 4.6.1 Tethering flies onto the dissection stage

A fly was cold anesthetized for 5 min. Then it was positioned onto the underside of a dissection stage and its wings were removed near their base using forceps. The thorax was then pressed through a hole (Etchit, Buffalo, MN) in the stage’s steel shim (McMaster-Carr, USA; 0.001” Stainless Steel, type 316 soft annealed; Part #2317K11). Afterwards, the stage was turned upside down and a tiny drop of UV-curable glue (Bondic, Aurora, ON Canada) was placed onto the scutellum, to fix the fly in place.

#### 4.6.2 Opening the thoracic cuticle

The stage was filled with saline solution **(Table 2)**. A 30 G syringe needle was then used to cut a small rectangular hole (smaller than the 600 μm diameter window) into the dorsal thoracic cuticle. The hole was made by inserting the needle into the posterior thorax close to the scutellum. Then three lines were cut into the lateral and anterior thorax. A final line was cut to complete a rectangular opening. The resulting piece of cuticle was then removed using forceps.

#### 4.6.3 Clearing out thoracic tissues

Residual degraded IFMs were removed from the opened thorax using forceps. Then, a pulled (P-1000, Sutter instrument, USA) glass needle (30-0018, Harvard Apparatus, USA) was used to detach small tracheal links between a large piece of trachea and the left side of the gut. The left salivary gland was then also removed using forceps.

#### 4.6.4 Displacing thoracic organs using the manipulator arm

The manipulator arm was positioned on top of the stage with its tip visible. The dissection stage was positioned with the fly’s head pointing toward the experimenter. The arm tip was then inserted into the thorax using a 3-axis manipulator (DT12XYZ, ThorLabs, USA). The tip of the arm was then inserted to the (experimenter’s) right side of the gut near the middle of the proventriculus. The tip was inserted deep enough to be below the crop and salivary glands but not to touch the VNC. Once the tip of the arm was on the right side of the salivary gland, crop, and gut, it was pulled towards the left side of the thoracic cavity, making a space for the closed implant.

#### 4.6.5 Positioning the implant

Once the flies’ organs were held securely onto the left side of the thoracic cavity by the manipulation arm, the implant was closed in the air using forceps and then transferred into the saline solution filling the dissection stage. The closed implant was then positioned in front of the fly on the stage. A thinner pair of forceps was next used to insert the implant into the animal’s thorax. Finally, a glass needle was used to adjust the location of the implant and to keep it at the appropriate height, allowing it to open passively. Once open, the glass needle was used to gently press the left side of the implant towards the bottom of the thorax while the arm was removed, and to remove any bubbles on the implant.

#### 4.6.6 Sealing the thoracic hole with a numbered, transparent window

Once the implant was well positioned, a syringe needle (15391557, Fisher Scientific, USA) was used to remove saline solution from the stage. A window was then positioned on top of the cuticular hole and centered with the identification number on the posterior of the thorax, near the scutellum. A wire was then used to add tiny drops of UV curable glue between the window and the surrounding thoracic cuticle, beginning from the right side of the scutellum and finishing on the left side. Saline solution was then added back to the stage. The cured UV glue, previously tethering the fly to the stage, was removed using a needle. The saline solution was then also removed and the window was fully sealed by placing and curing UV glue onto the fly’s posterior cuticle near the scutellum.

#### 4.6.7 Dismounting flies from the dissection stage

Once the thoracic hole was fully sealed by a transparent window, the fly was dismounted from the dissection stage by gently pushing the front of the thorax through the hole in the steel shim. The fly was then returned to a vial of food to recover.

### 4.7 *Drosophila melanogaster* experiments

All flies were raised on standard food on a 12h light:12h dark cycle. Experiments for each particular study were performed at a consistent time of day to exclude the possibility of circadian-related confounding factors.

### 4.8 Long-term study of survival and locomotion

Female flies expressing CsChrimson in Moonwalker Descending Neurons (MDNs) [22](*UAS-CsChrimson / Act88F-Rpr; VT50660*.*p65AD(attp40) /+; VT44845*.*Gal4DBD(attp2) /+*)**(Figure 2)** were implanted at five days-post-eclosion (dpe). For this experiment, before implantation, implants were dipped in a 30 mg/ml dextran solution (#31392, Sigma-Aldrich, Switzerland) while mechanically closed. Implants were then taken out of the solution and dried using a twisted Kimwipe (5511, Kimberly-Clark, USA). This step was performed to fix implants in a closed position. However, we later discovered that dextran is not required to close implants and we removed this step. Implants were then positioned in the fly’s thorax as described above. The number of days following implantation is denoted as ‘days-post-implantation’ (dpi). Age and gender-matched control animals were selected from the same parental cross. For longevity studies, flies were housed individually in food vials and assessed every 1-2 days.

Studies of locomotion were performed at 1-3 dpi, 14-16 dpi, and 28-30 dpi. Animals were individually cold-anaesthetized and then transferred to rounded square arenas for optogenetic activation and video recording. Each recording consisted of 30 s of spontaneously generated behaviors (primarily walking and grooming), followed by three 3 s periods of optogenetic stimulation at 590 nm (6 mW / cm^2^) with 10 s interstimulus intervals. Therefore, each recording session was 59 s long.

To process video data, flies’ centroids were tracked using a customized version of Tracktor [57]. Their orientations were then extracted using a neural network (implemented in PyTorch [58]) that was trained on hand-labeled data. The network consisted of two convolutional layers followed by three fully connected layers. All layers, except for the final one, were followed by a ReLU activation function [59]. We also applied dropout after the first two fully connected layers with 0.2 probability [60]. To train the network, we hand annotated a total of 300 samples in three orientations (head up, head down, and sideways). The grayscale images were then cropped using Tracktor centroid locations and resized to 32 *×* 32 pixels. During training, we randomly applied affine transformations (20 degrees of rotation, 5 pixels of translation, and 0.2 scaling factor), horizontal, and vertical flip augmentations with a 0.5 probability. We used PyTorch’s torchvision package for all data augmentation. The network was trained with cross-entropy loss using 80% of the data. We used an Adam optimizer with a learning rate of 0.001, without weight decay and learning rate drop [61]. We trained for 1000 epochs and selected the weights with the best test error.

Translational velocities were computed by applying a second order Savitzky-Golay filter with a first-order derivative to centroid positions. The sign for the velocity values was set to negative for movements counter to the animal’s heading direction. The ‘Backward walking response slope’ metric was calculated as the acceleration from the beginning of each stimulation period to the minimum velocity (maximum backward speed) reached on that period. The ‘Backward walking distance traveled’ metric was computed as the left Riemann sum of the velocity curves during each stimulation period. We only considered frames where the velocity was negative. Finally, the ‘Maximum negative translational velocity’ is the minimum velocity value reached on each stimulation period.

### 4.9 Long-term anatomical imaging of the VNC

Female flies expressing GFP throughout the nervous system (*Act88F-Rpr/+; GMR57C10-Gal4/UAS-GFP; +/+*) **(Figure 1H)** were implanted at 4-6 dpe and kept individually in food vials. At 1-3 dpi, 14-16 dpi, and 28-30 dpi, flies were tethered onto a remounting stage and 25 imaging volumes of 100 μm depth (1 μm stepsize) were acquired using a two-photon microscope (Bergamo II microscope, ThorLabs, USA) and a 930 nm laser (MaiTai DeepSee, Newport Spectra-Physics, USA) with 20 mW of power at the sample location. We acquired 0.1 volumes-per-second (vps) using a Galvo-Resonance scanner [17]. The 25 images per depth were then registered to one another using the HyperStackReg module in Fiji [62] and a rigid body transformation. These registered images were next projected along the time axis into one standard deviation image. The resulting volume was then depth color-coded using Fiji’s Temporal-Color macro.

### 4.10 Long-term functional recording of the VNC

A female fly expressing GCaMP6f and tdTomato throughout the nervous system (*Act88F-Rpr/+; GMR57C10-Gal4/UAS-GCaMP6f; UAS-tdTomato/+*) was implanted at 5 dpe. The same fly was then mounted onto the two-photon imaging stage at 1, 5 and 10 dpi. One horizontal imaging plane of the prothoracic neuromere was acquired using a two-photon microscope at 930 nm with 25 mW of power. Three horizontal z-plane images were acquired using a Galvo-Resonance scanner and averaged into one frame at an imaging rate of 10.7 fps. Behavior frames were acquired simultaneously (as in [17]) at a rate of 80 fps.

### 4.11 Long-term study of chordotonal organ degradation in the VNC following leg amputation

Female flies expressing GFP in their chordotonal organs (*Act88F-Rpr/+; iav-Gal4/UAS-GFP; +/+*) **(Figure 3)** were implanted at 1 dpe. A z-stack of the VNC was recorded at 1 dpi, using a two-photon microscope at 930 nm with 55 mW of laser power. Flies were anesthetized with carbon dioxide (1.8l/min) supplied ventrally while recording z-stacks. Z-stacks consisted of 576×384 pixel frames taken every 1 μm over a total depth of 100 μm (i.e., 100 frames per volume). The front left leg was then removed at the thorax-coxa joint using dissection scissors (#15300-00, Fine Science Tools, Germany). A second z-stack was then immediately recorded. Flies were kept individually in food vials and imaged every day using the same recording parameters until 15 dpi. Fiji’s linear stack alignment with the SIFT registration plugin [63] was then used to register all the projected z-stacks to the first z-stack. A custom python script was then used to draw and extract the mean fluorescence of specific regions of interest. Mean fluorescence within these regions were measured for each day and normalized across animals by dividing them by the mean fluorescence on the first day.

Flies’ nervous systems were dissected and fixed with paraformaldehyde (441244, Sigma-Aldrich, USA) at 20 dpi. Samples were then stained for nc82 as in [17]. This allowed us to acquire confocal images that included both neuropil landmarks and endogenous GFP expression. Confocal laser intensities and PMT gains were manually selected to avoid pixel saturation. These confocal z-stacks were then projected into 2D using Fiji’s standard deviation projection. The standard deviation projection of GFP expression is shown as an inverted image **(Figure 3D,E)**. A custom python script was written to detect the VNC’s boundaries using the standard deviation projection of nc82 images. This contour was detected using the Open CV library and then drawn onto GFP standard deviation projection images.

### 4.12 Recording neural population activity before, during, and after feeding

Female flies (5 dpe) expressing a calcium indicator, GCaMP6f, and an anatomical marker, tdTomato, throughout the nervous system (*Act88F-Rpr/+; GMR57C10-Gal4/UAS-GCaMP6f; UAS-tdTomato/+*) **(Figure 4)** were starved for 21-23 h on a wet Kimwipe (5511, Kimberly-Clark, USA). They were then implanted without a thoracic window, and kept on the dissection stage (the remounting stage was not used here) to limit the number of interventions. Animals were then positioned under a two-photon microscope where they could walk on a spherical treadmill consisting of an air-supported (0.8 L/min) foam ball (Last-A-Foam FR7106, General Plastics, USA) with a diameter of 1cm [17]. Coronal cross-sections of the cervical connectives were then imaged at 930 nm with a laser power of 15 mW. We achieved a 16 frames-per-second (fps) imaging rate by using a Galvo-Resonance scanner. In parallel, the behavior of the flies was recorded using seven cameras at 80 fps. Ball rotations were also measured along three axes using two optic flow sensors [11, 17]. We recorded neural activity and behavior in trials of approximately four minutes each. First, four trials were recorded. Then, the foam ball was lowered and recording continued while flies fed on a solution consisting of either (i) 1 ml deionized water, 8 mg of sucrose (A2188.1000, Axon Lab, Switzerland) and 1 mg of Amaranth dye (A1016, Sigma-Aldrich, USA), (ii) a low concentration caffeine solution consisting of 1 ml deionized water, 8 mg caffeine (C0750, Sigma-Aldrich, USA), 8 mg of sucrose and 1mg of Amaranth, or (iii) a high concentration supersaturated caffeine solution consisting of 1 ml deionized water, 40 mg caffeine, 8 mg of sucrose and 1mg of Amaranth. Animals were fed using a pulled glass needle (P-1000, Sutter instrument, USA; puller parameters-Heat: 502; Pull:30; Velocity: 120; Time: 200; Pressure: 200). A tiny drop of UV curable glue (Bondic, Aurora, ON Canada) was added near the tip of the needle to prevent the solution from travelling up on the needle. The needle was positioned in front of the flies using a manipulator (uMp-3, Sensapex, Finland). After feeding, the spherical treadmill was repositioned below the fly and eight more imaging trials were acquired.

#### 4.12.1 Motion correction of two-photon imaging data

We used custom Python code unless otherwise indicated. For all image analysis, the y-axis is ventral-dorsal along the fly’s body, and the x-axis is medial-lateral. Image and filter kernel sizes are specified as (y, x) in units of pixels. Recordings from the thoracic cervical connective suffer from large inter-frame motion including large translations, as well as smaller, non-affine deformations. Because calcium indicators (e.g., GCaMP6f) are designed to have low baseline fluorescence, they are challenging to use for motion correction. Therefore, we relied on signals from the co-expressed red fluorescent protein, tdTomato, to register both the red (tdTomato) and the green (GCaMP6f) PMT channel images. First, we performed center-of-mass (COM) registration of each recorded frame to remove large translations and cropped the background regions around the neck connective (from 480×736 to 352×576). Then, we computed the motion field of each red frame relative to the first recorded frame using optic flow and corrected both red and green frames for the motion using bi-linear interpolation. The algorithm for optic flow motion correction was previously described in [17]. We only used the optic flow component to compute the motion fields and omitted the feature matching constraint. We regularized the gradient of the motion field to promote smoothness (*λ* = 800). Python code for the optic flow motion correction (ofco) package can be found at https://github.com/NeLy-EPFL/ofco.

#### 4.12.2 Correction for uneven illumination

We observed that absolute fluorescence values were slightly lower on the right side of the connective than the left side, likely due to scattering by thoracic organs that are pushed to the right by the implant. To correct for this uneven absolute fluorescence, we computed the mean of all motion corrected frames across time. We then median filtered and low-pass filtered the resulting image (median filter: (71,91), Gaussian filter: *σ* = 3) to remove the features of individual neurons and retain only global, spatial changes in fluorescence. We then computed the mean across the y axis to obtain a fluorescence profile in the x (left - right) axis and fit a straight line to the most central 200 pixels. To correct for the decrease in fluorescence towards the right side, we multiplied the fluorescence with the inverse value of this straight line fit to the x-axis profile. Note that this correction only aids in the visualisation of fluorescence, and does not have any impact on the computation of Δ*F/F* because, for a given pixel, both the fluorescence at each time point, and its baseline fluorescence are multiplied by the same constant factor.

#### 4.12.3 Denoising calcium imaging data

To denoise registered and corrected data, we used an adapted version of the DeepInterpolation algorithm [64]. Briefly, DeepInterpolation uses a neuronal network to denoise a microscopy image by “interpolating” it from temporally adjacent frames. A U-Net is trained in an unsupervised manner using 30 frames (around 2s) before and 30 frames after the target frame as an input and the current frame as an output. Thus, independent noise is removed from the image and components that dynamically evolve across time are retained. We modified the training procedure to fit one batch into the 11GB RAM of a Nvidia GTX 2080TI graphics card: rather than use the entire frame (352×576 pixels), we used a subset of the image (352×288 pixels) during training. We randomly selected the x coordinate of the subset. During inference, we used the entire image. We verified that using different images sizes during training and inference did not change the resulting denoised image outside of border regions. We trained one model for each fly using 2000 randomly selected frames from one of the trials before feeding and applied it to all of subsequent frames. Training parameters are outlined in Table 1. The adapted DeepInterpolation algorithm can be found on the “adapttoR57C10” branch of the following GitHub repository: https://github.com/NeLy-EPFL/deepinterpolation

**Table 1:** Training parameters for DeepInterpolation.

| Parameter | Value |
| --- | --- |
| Number of training frames | 2000 |
| Number of frames pre/post current frames | 30 |
| Omission of frames pre/post current frame | 0 |
| Number of iterations through training data | 1 |
| Learning rate | 0.0001 |
| Learning decay | 0 |
| Batch size | 4 |
| Steps per epoch | 5 |
| Number of GPUs | 1 |
| Number of workers | 16 |

#### 4.12.4 Generating Δ*F*/*F* videos

We show fluorescence values as Δ*F*/*F* **(Videos 7-10)**. This was computed as 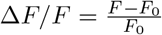, where *F* is the time varying fluorescence and *F*_0_ is the pixel-wise fluorescence baseline. To compute *F*_0_, we applied a spatial Gaussian filter (*σ* = 10) to images and convolved each pixel with a temporal window of 10 samples (around 0.6s). We then identified the minimum fluorescence of each pixel across all trials.

#### 4.12.5 Optic flow processing and classification of stationary periods

Optic flow sensors have been used to measure spherical treadmill rotations [11, 17] but they are inherently noisy. Therefore, we computed the moving average across 80 samples (around 200ms). From preprocessed sensor values, we computed the forward, sideways and turning velocities [11]. We classified stationary periods (no movements of the ball) as when the absolute values of each of the three ball rotation velocities are below a threshold optic flow value of 0.31m s^*−*1^ ≙ 0.01 rotations*/*s and at least 75% of the frames within the time ±0.5s of the sample are below this threshold. The latter criterion ensures that short stationary periods between bouts of walking would be excluded.

#### 4.12.6 Synchronisation of two-photon, optic flow, and camera data

We recorded three different data modalities at three different sampling frequencies: two-photon imaging data was recorded at approximately 16Hz, behavioral images from seven cameras were acquired at 80Hz, and ball movements using two optical flow sensors were measured at nearly 400Hz. Therefore, to synchronise these measurements for further analysis, we down-sampled all measurements to the two-photon imaging frame rate by averaging all behavioral and ball rotation samples acquired during one two-photon frame.

#### 4.12.7 Data analysis for caffeine ingestion experiment

To compute Δ*F/F* traces for each trial—as shown in Figure 4—we averaged the fluorescence across the entire cervical connective and computed the Δ*F/F* of this time series as for individual pixels (see above). To analyze the temporal progression of fluorescence waves, we first identified the time of peak fluorescence across the entire cervical connective *T*_*peak*_. All times are given relative to the time of that peak. We then computed the mean fluorescence across time within manually selected regions of interest (dorsal, lateral, and ventral connective, as well as giant fiber neurons) and represent them normalised to their minimum and maximum values. We smoothed the time series with a Gaussian filter (*σ* = 3 ≙0.18s). To identify the peak time for each pixel, we applied a temporal Gaussian filter (*σ* = 10 ≙0.62s) and spatial Gaussian filter (*σ* = 1) and searched for the maximum fluorescence value within *T*_*peak*_ ± 10s. In Figure 4 we show the mean fluorescence during periods when the fly was stationary (i.e., not moving the ball).

## 5 Supplementary Tables

**Table 2:** Saline solution

| Chemical | mM |
| --- | --- |
| NaCl | 103 |
| KCl | 3 |
| NaHCO<sub>3</sub> | 26 |
| NaH<sub>2</sub>PO<sub>4</sub> | 1 |
| CaCl<sub>2</sub> (1M) | 4 |
| MgCl<sub>2</sub> (1M) | 4 |
| Trehalose | 10 |
| TES | 5 |
| Glucose | 10 |
| Sucrose | 2 |

**Table 3:**
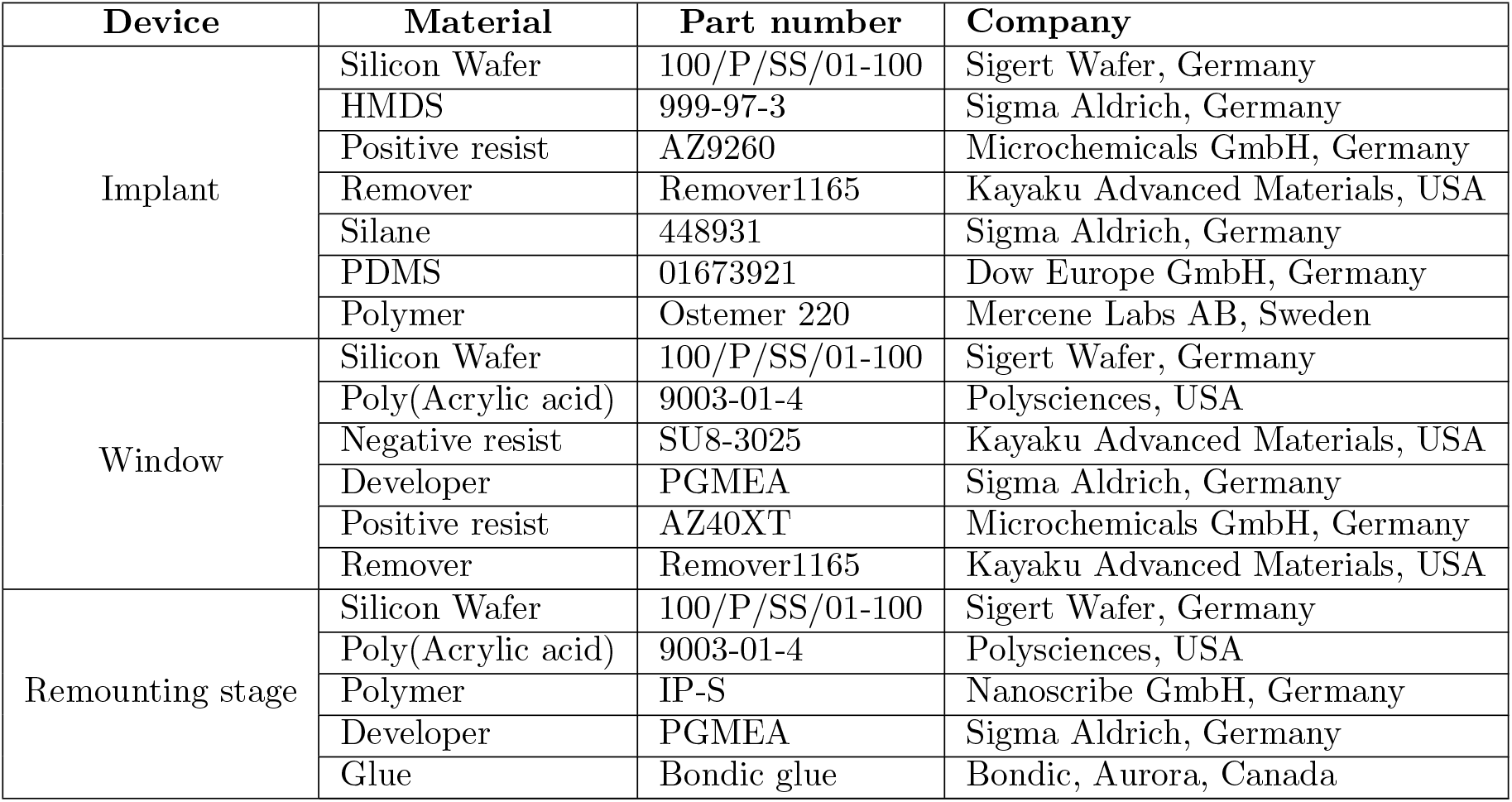
Main materials for long-term imaging tool fabrication

| Device | Material | Part number | Company |
| --- | --- | --- | --- |
| Implant | Silicon Wafer | 100/P/SS/01-100 | Sigert Wafer, Germany |
|  | HMDS | 999-97-3 | Sigma Aldrich, Germany |
|  | Positive resist | AZ9260 | Microchemicals GmbH, Germany |
|  | Remover | Remover1165 | Kayaku Advanced Materials, USA |
|  | Silane | 448931 | Sigma Aldrich, Germany |
|  | PDMS | 01673921 | Dow Europe GmbH, Germany |
|  | Polymer | Ostemer 220 | Mercene Labs AB, Sweden |
| Window | Silicon Wafer | 100/P/SS/01-100 | Sigert Wafer, Germany |
|  | Poly(Acrylic acid) | 9003-01-4 | Polysciences, USA |
|  | Negative resist | SU8-3025 | Kayaku Advanced Materials, USA |
|  | Developer | PGMEA | Sigma Aldrich, Germany |
|  | Positive resist | AZ40XT | Microchemicals GmbH, Germany |
|  | Remover | Remover1165 | Kayaku Advanced Materials, USA |
| Remounting stage | Silicon Wafer | 100/P/SS/01-100 | Sigert Wafer, Germany |
|  | Poly(Acrylic acid) | 9003-01-4 | Polysciences, USA |
|  | Polymer | IP-S | Nanoscribe GmbH, Germany |
|  | Developer | PGMEA | Sigma Aldrich, Germany |
|  | Glue | Bondic glue | Bondic, Aurora, Canada |

**Table 4:**
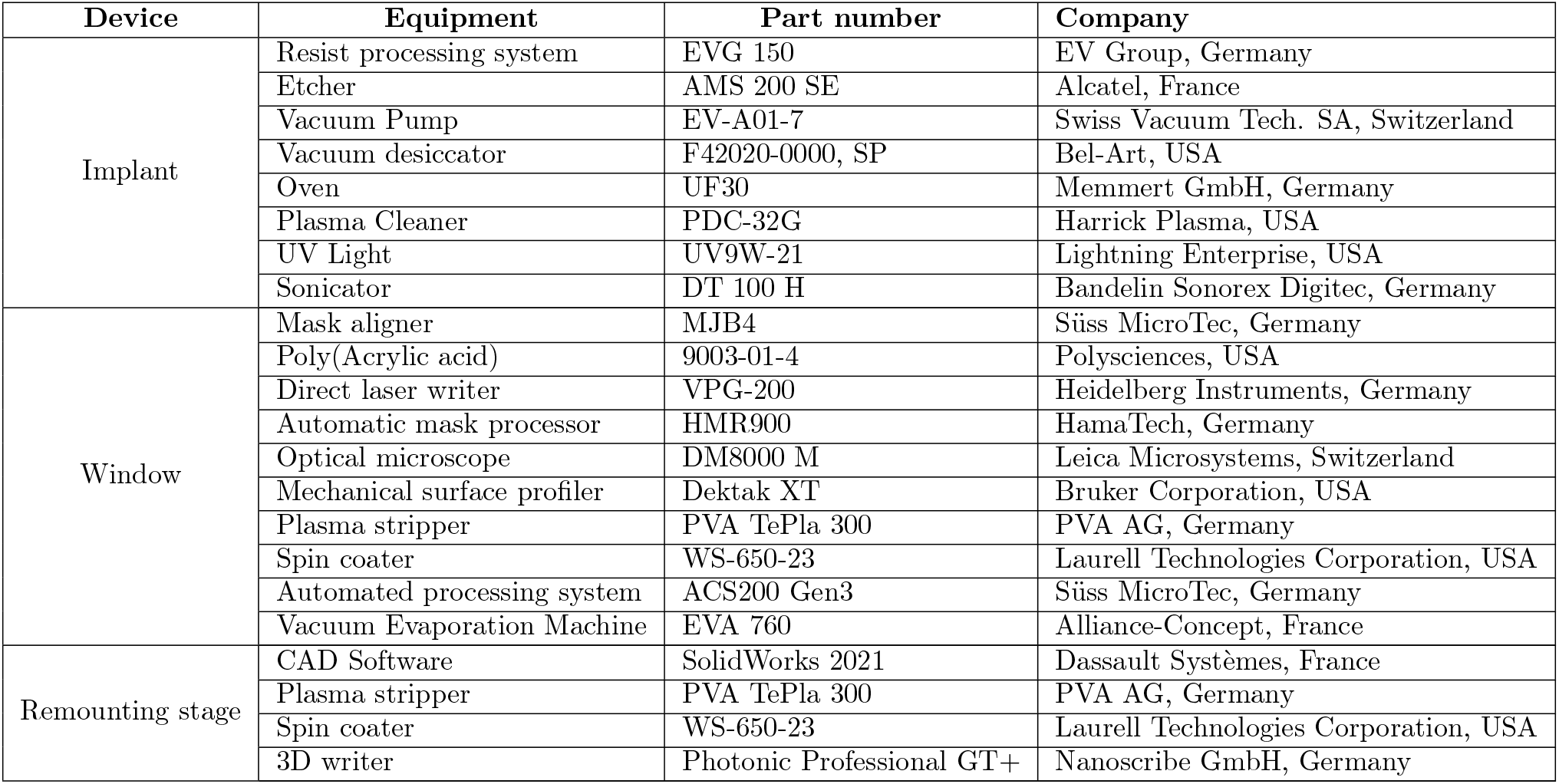
Main equipment for long-term imaging tool fabrication

## 6 Supplementary Figures

**Figure S1:**
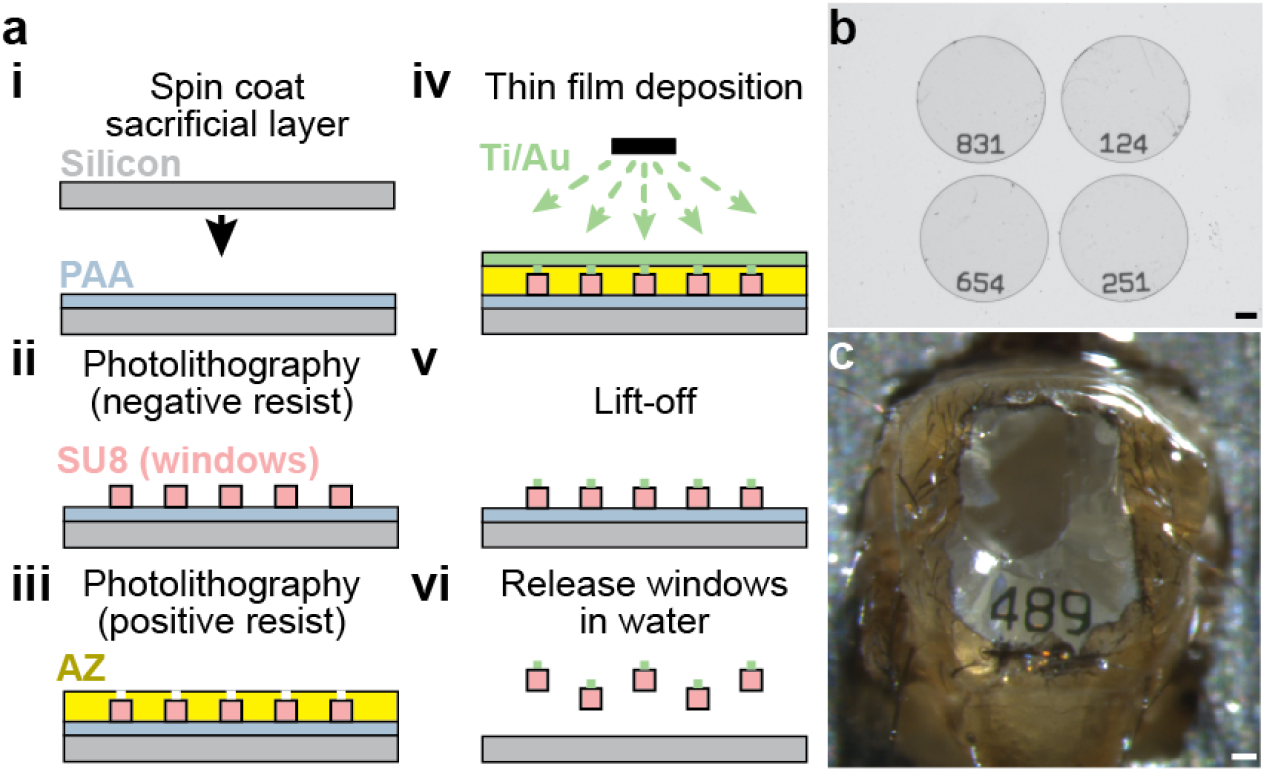
Fabrication of number-coded and optically transparent thoracic windows. **(A)** Thoracic windows are fabricated by performing the following steps. **(i)** A sacrificial layer of PAA is spin-coated onto a silicon wafer, using photolithography. **(ii)** SU-8 windows are structured onto the sacrificial layer. **(iii)** A positive resist, AZ, is cross-linked to mark number openings. **(iv)** Ti/Au is vapor deposited. **(v)** The AZ layer is lifted off. **(vi)** Finally, the numbered windows are released in water. **(B)** This process yields transparent SU-8 windows with thin Ti/Au numbers. Scale bar is 100 μm. **(C)** A window on an implanted animal, permitting a view of thoracic organs and tracking of this animal’s identity. Scale bar is 50 μm.

**Figure S2:**
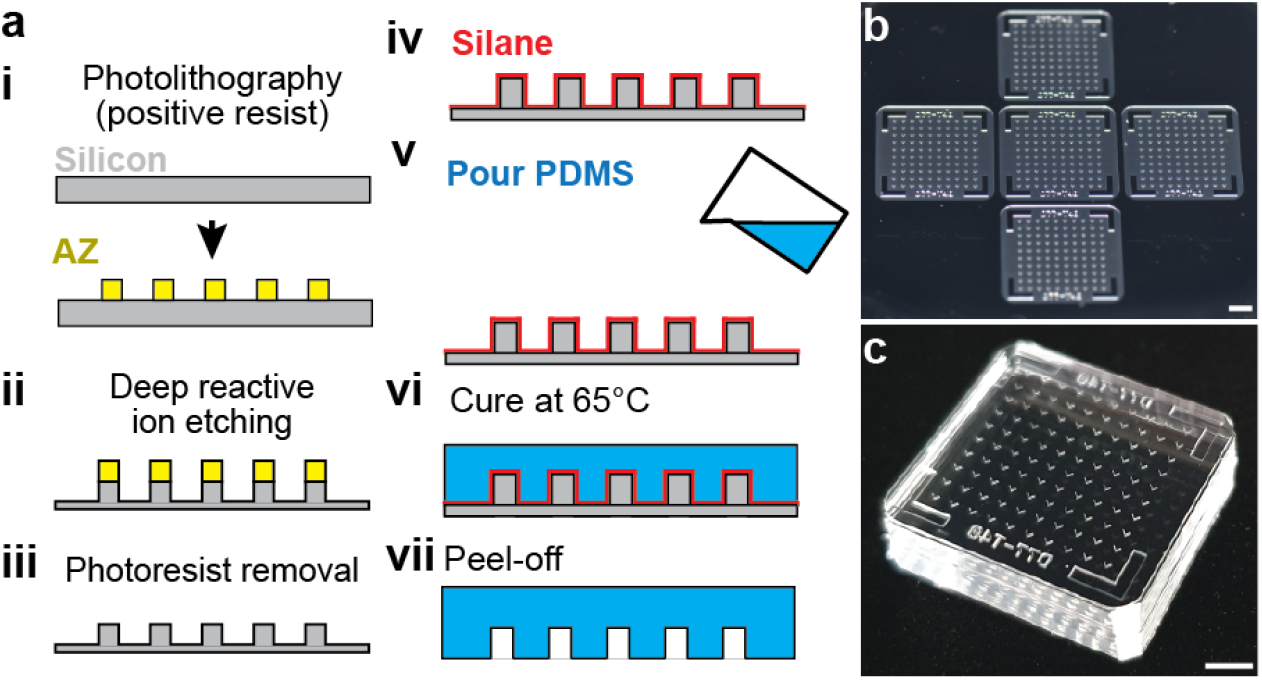
Fabrication of molds used to cast implants. **(A)** Implant molds are fabricated by performing the following steps. **(i)** Through photolithography, a positive resist, AZ, is cross-linked onto a silicon wafer to form a temporary mask. **(ii)** Deep reactive ion etching is used to sculpt the silicon wafer. **(iii)** The photoresist is removed. **(iv)** Subsequently, this silicon piece is silanized. **(v)** PDMS is then poured, **(vi)** cured, and **(vii)** peeled off. **(B)** This process yields a single large piece. Scale bar is 0.5 cm. **(C)** This large piece is cut into five individual implant molds. Scale bar is 0.5 cm.

**Figure S3:**
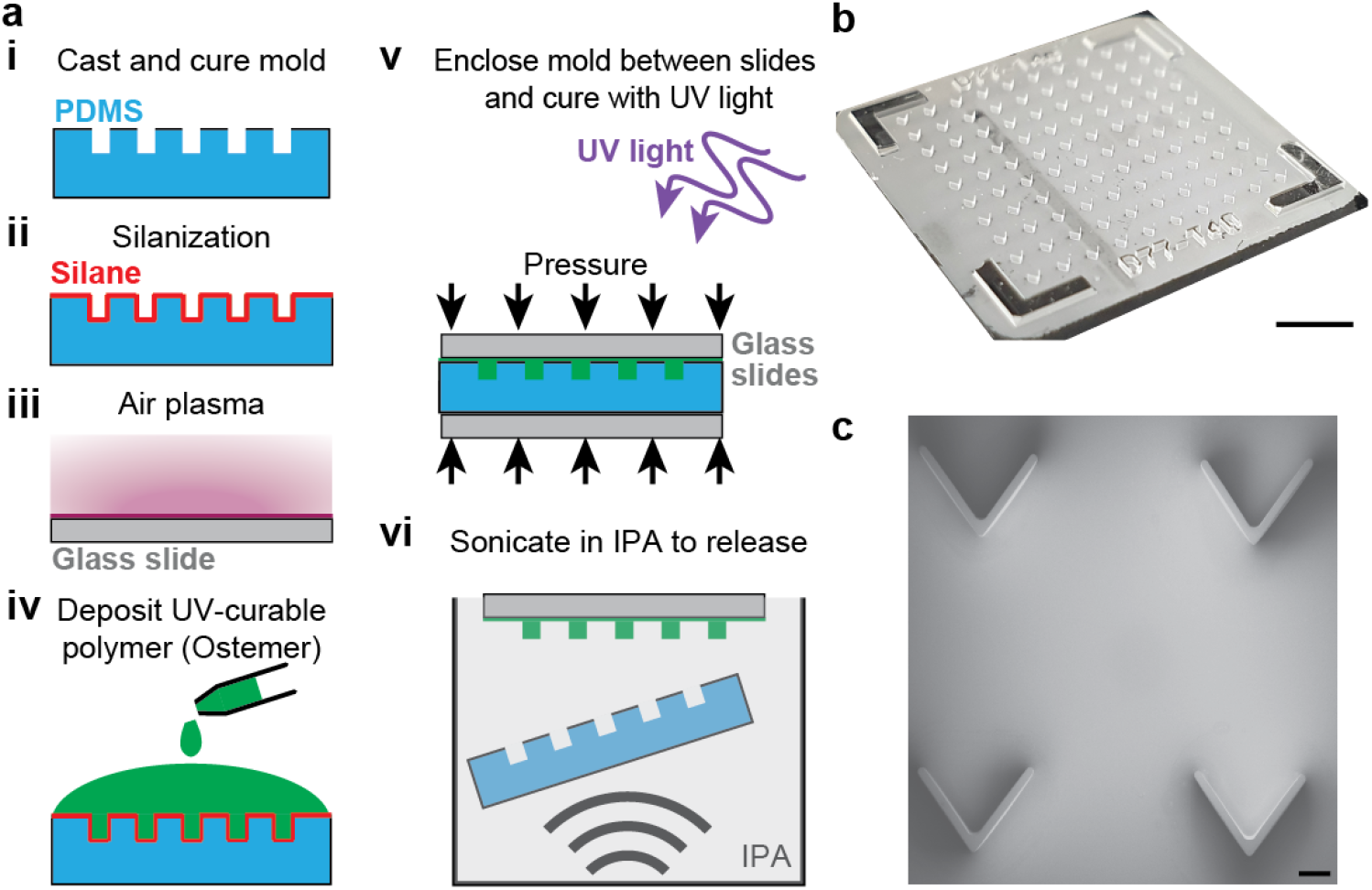
Fabrication of implants. **(A)** Implants are fabricated by performing the following steps. **(i)** PDMS molds are cast, cured, and cut into pieces. **(ii)** PDMS molds are silanized. **(iii)** A glass slide is plasma treated to promote adhesion. **(iv)** A UV curable polymer is deposited onto the PDMS mold. **(v)** This composite is sandwiched between glass slides and exposed to UV light. **(vi)** The mold is sonicated to release in IPA. **(B)** This high-throughput process yields 100 implants in a single mold. Scale bar is 0.5 cm. **(C)** A scanning electron microscopy image confirms the precision of implant fabrication. Scale bar is 200 μm.

**Figure S4:**
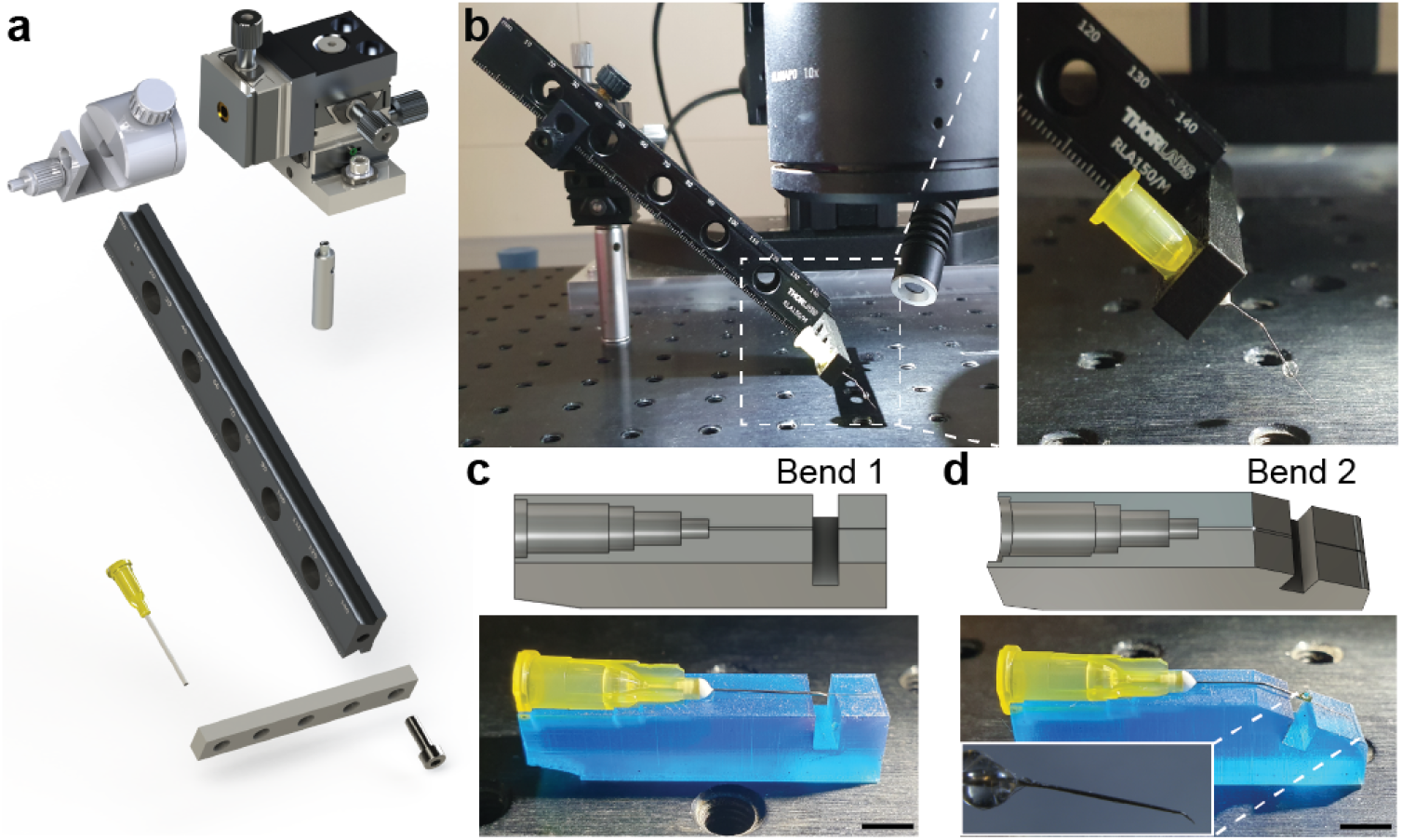
Fabrication of a manipulator arm to temporarily displace thoracic organs. **(A)** Exploded view of the manipulator arm and its component parts. **(B) (Left)** View of the manipulator arm mounted near the dissection microscope. **(Right)** Zoomed in view of the inset (dashed white lines) highlighting the bent needle tip. **(C)** 3D printed piece used to guide gluing of the pin to the syringe needle. Scale bar is 0.5 cm. **(D)** 3D printed piece used to guide bending the manipulator arm tip. Inset shows a zoomed-in view of the arm’s tip.

**Figure S5:**
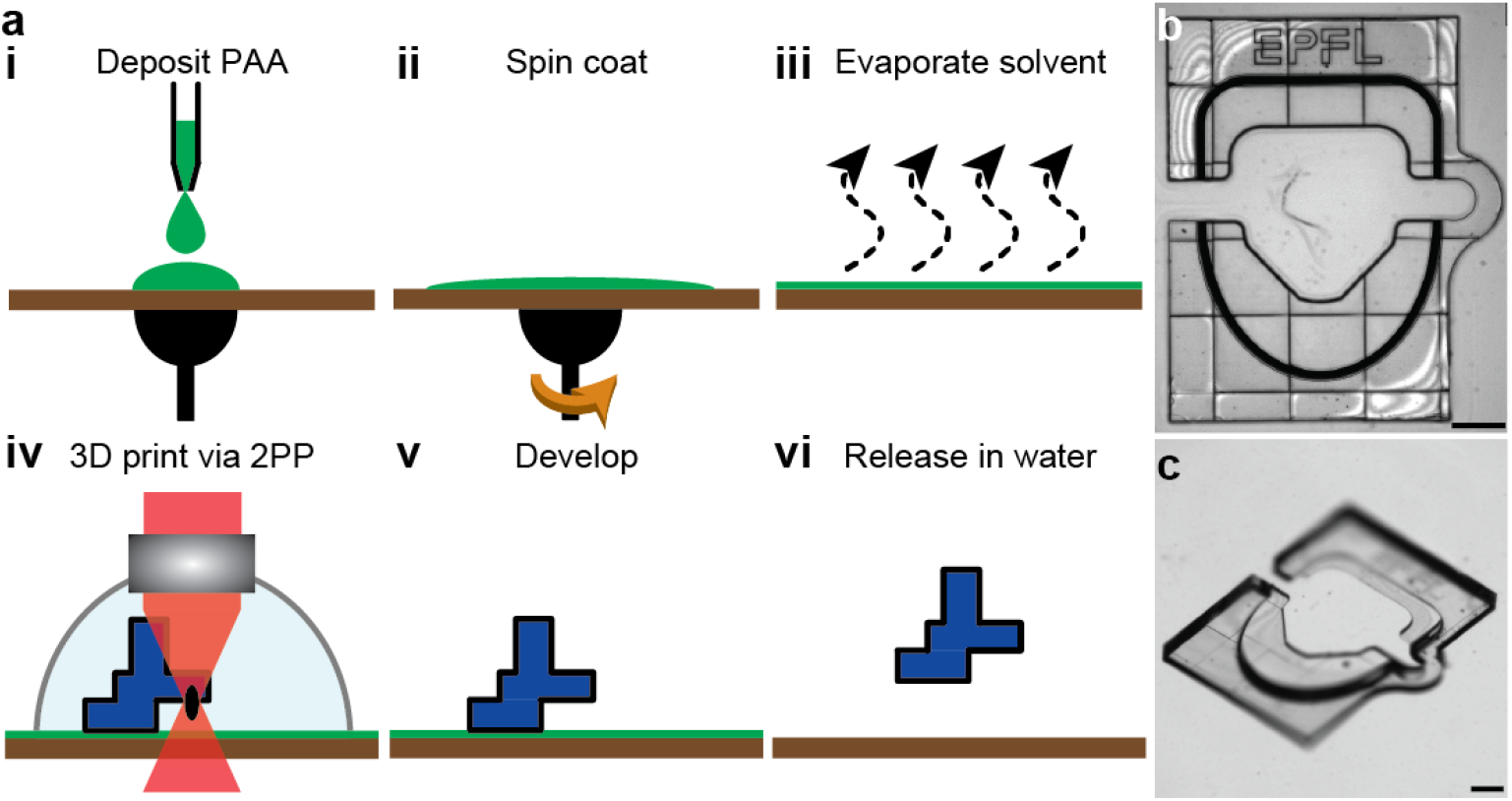
Fabrication of remounting stage. **(A)** A water soluble sacrificial solution is **(i)** deposited and **(ii)** spin-coated to ensure a thin layer. **(iii)** The water in the solution is evaporated, leaving a dry water soluble layer. **(iv)** The remounting stage is 3D printed using two-photon polymerization (2PP). **(v)** This is followed by development in a solvent. **(vi)** Finally, the piece is released in water. **(B)** A microscope image of the remounting stage before releasing it in water. Scale bar is 0.25 mm. **(C)** Another view of the remounting stage illustrating its ergonomic design for fly tethering. Scale bar is 0.25 mm

**Figure S6:**
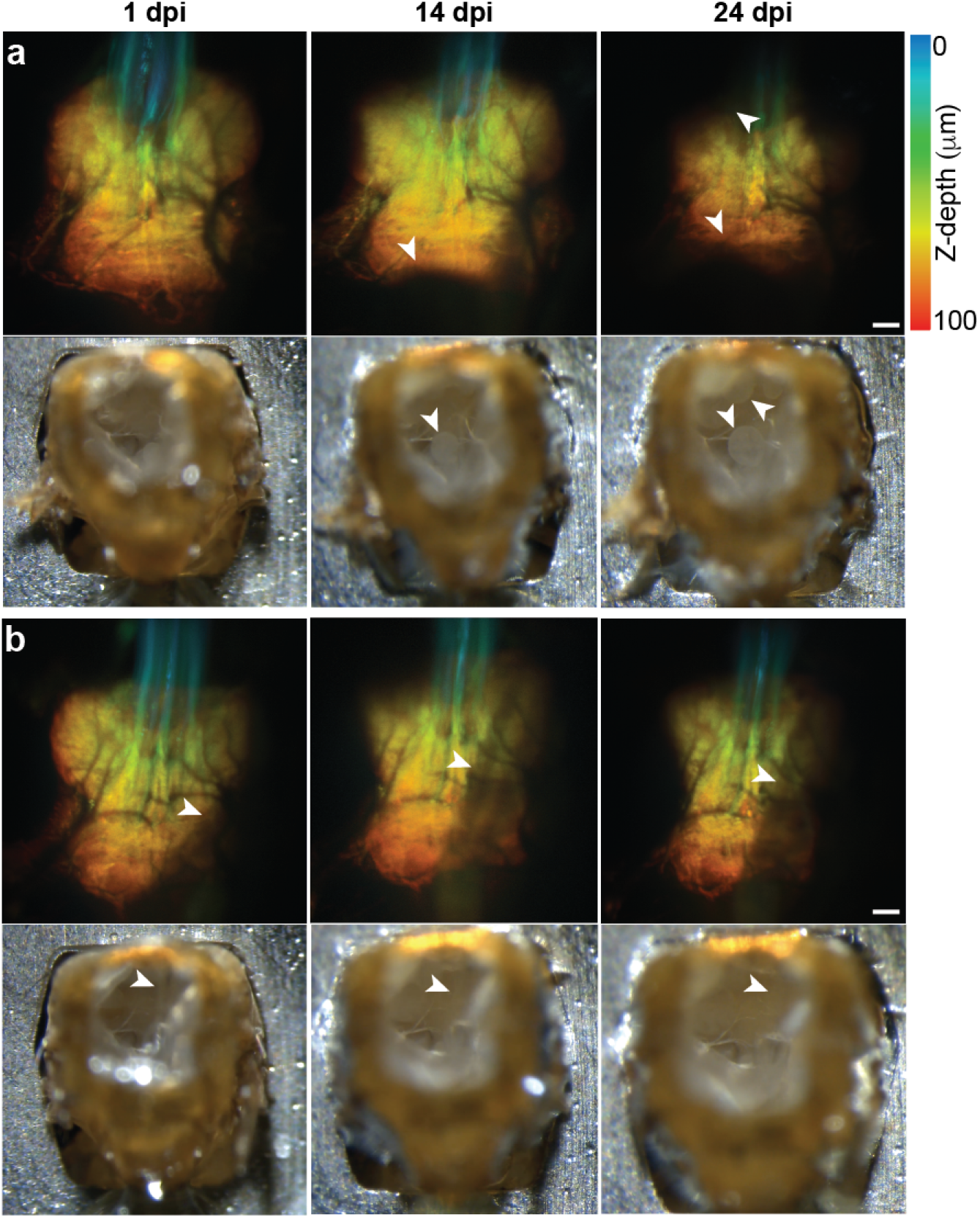
Potential organ movements within the thorax after implantation. Following two implanted animals at **(left)** 1 dpi, **(middle)** 14 dpi, and **(right)** 24 dpi. Shown are two animals with image-obscuring movements of the **(A)** fat bodies, or **(B)** salivary glands. **(top row)** Two-photon images of the animal’s VNC expressing GFP throughout the nervous system. White arrowheads indicate **(A)** fat bodies or **(B)** salivary glands shifting over time leading to an obscured view of the VNC. Z-stacks are depth color-coded (100 μm). Scale bar is 25 μm. **(bottom row)** The same animal’s dorsal thorax, visualized using a dissection microscope.

**Figure S7:**
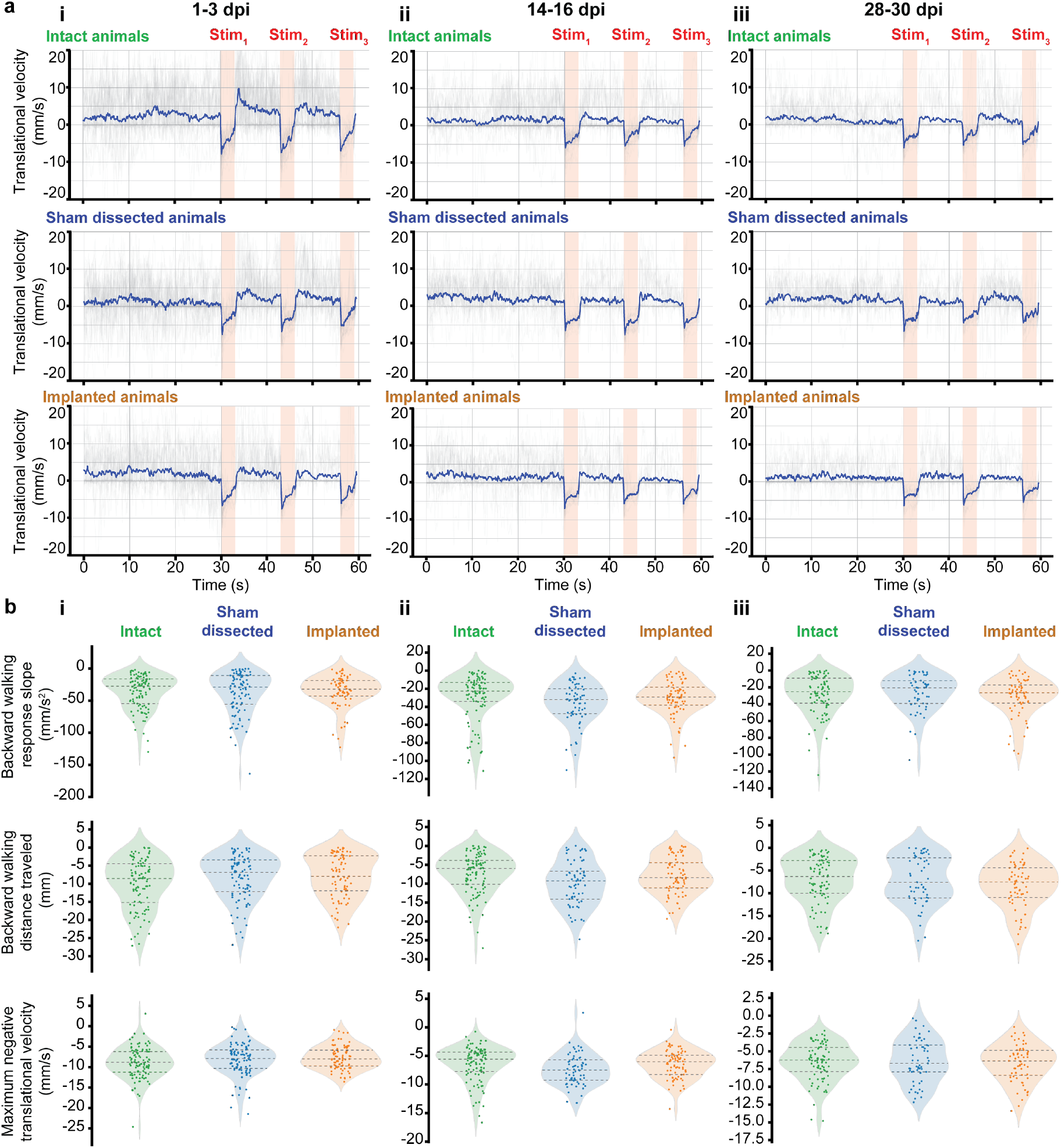
Impact of implantation and windows on behavior, separated by age post implantation. **(A)** Translational velocities of intact **(top)**, sham dissected **(middle)**, and implanted **(bottom)** animals during 30 s of spontaneous behavior, followed by three optogenetic stimulation periods of 3 s each (pink, ‘Stim’). Shown are the raw (grey) and mean (blue) traces arranged by age: **(i)** 1-3 dpi, **(ii)** 14-16 dpi, or **(iii)** 28-30 dpi. **(B)** From these time-series data, summary statistics include **(top)** the initial negative slope in translational velocity—backward walking—upon optogenetic stimulation, **(middle)** the integrated translational velocity over the entire optogenetic stimulation period, and **(bottom)** the peak negative translational velocity over the entire optogenetic stimulation period. Data are sorted by age as in panel A. A Kruskal-Wallis statistical test was used to compare behaviors across the three groups. A posthoc Conover’s test with a Holm correction was used to perform pairwise comparisons. Significant differences were found only at 14-16 dpi between the ‘Sham dissected’ and two other groups. Specifically, the ‘Sham dissected’ group showed significant differences (i) to the ‘Intact’ group for the backward walking response slope (*P* =0.03), (ii) to the ‘Intact’ group in the backwawrd walking distance traveled (*P* =0.01), and (iii) to the ‘Intact’ (*P* =0.004) and ‘Implanted’ groups (*P* =0.03) for the maximum negative translational velocity. No significant differences were observed between the ‘Intact’ and ‘Implanted’ groups.

**Figure S8:**
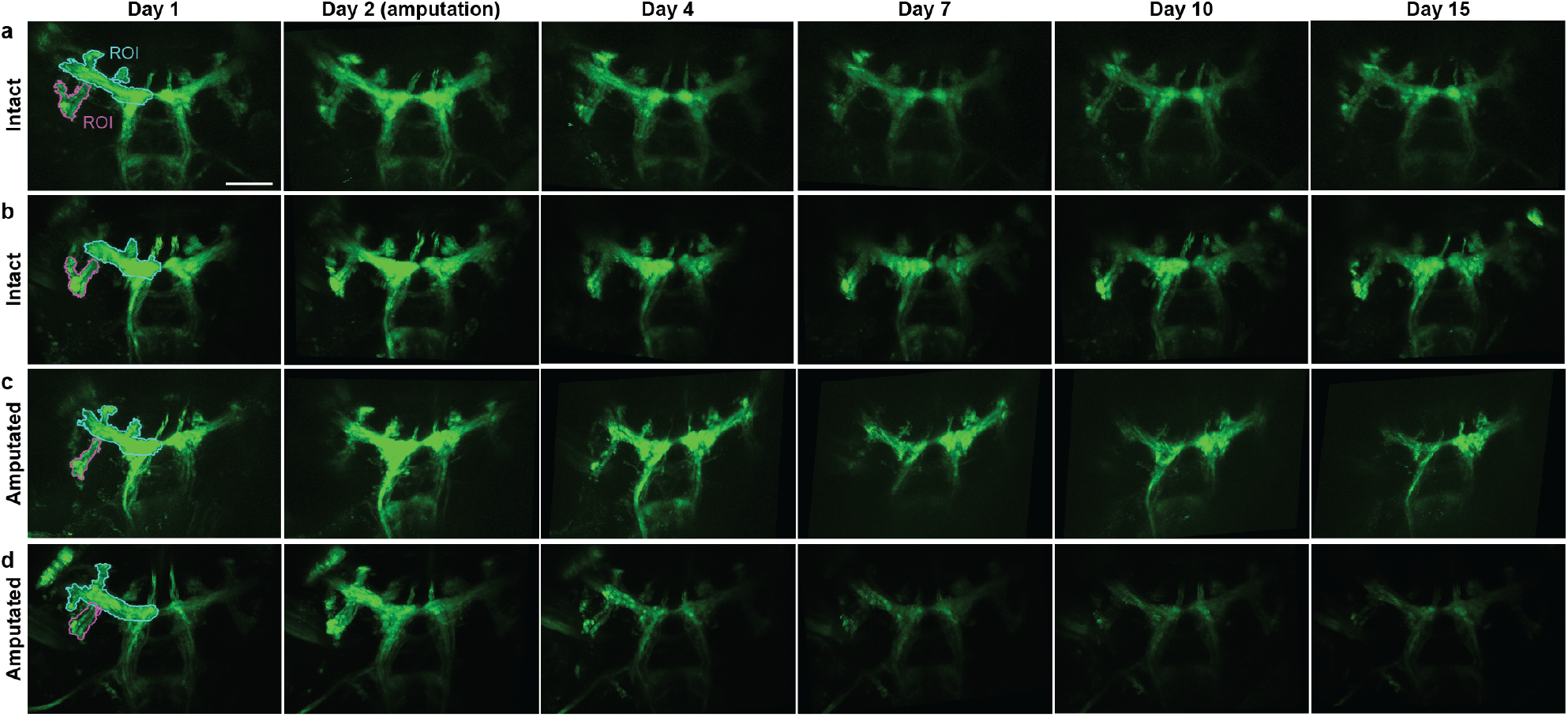
Long-term imaging of front leg chordotonal organs axon terminals in the VNC in intact or amputee animals. Maximum intensity projections of z-stacks taken at 1, 2, 4, 7, 10 and 15 dpi. Data are registered to images at 1 dpi. Scale bar is 50μm. Cyan and pink ROIs used for quantification in Figure 3 are indicated. Data are for **(A,B)** two control animals with intact legs and **(C,D)** two animals whose front left legs were amputated at 2 dpi.

**Figure S9:**
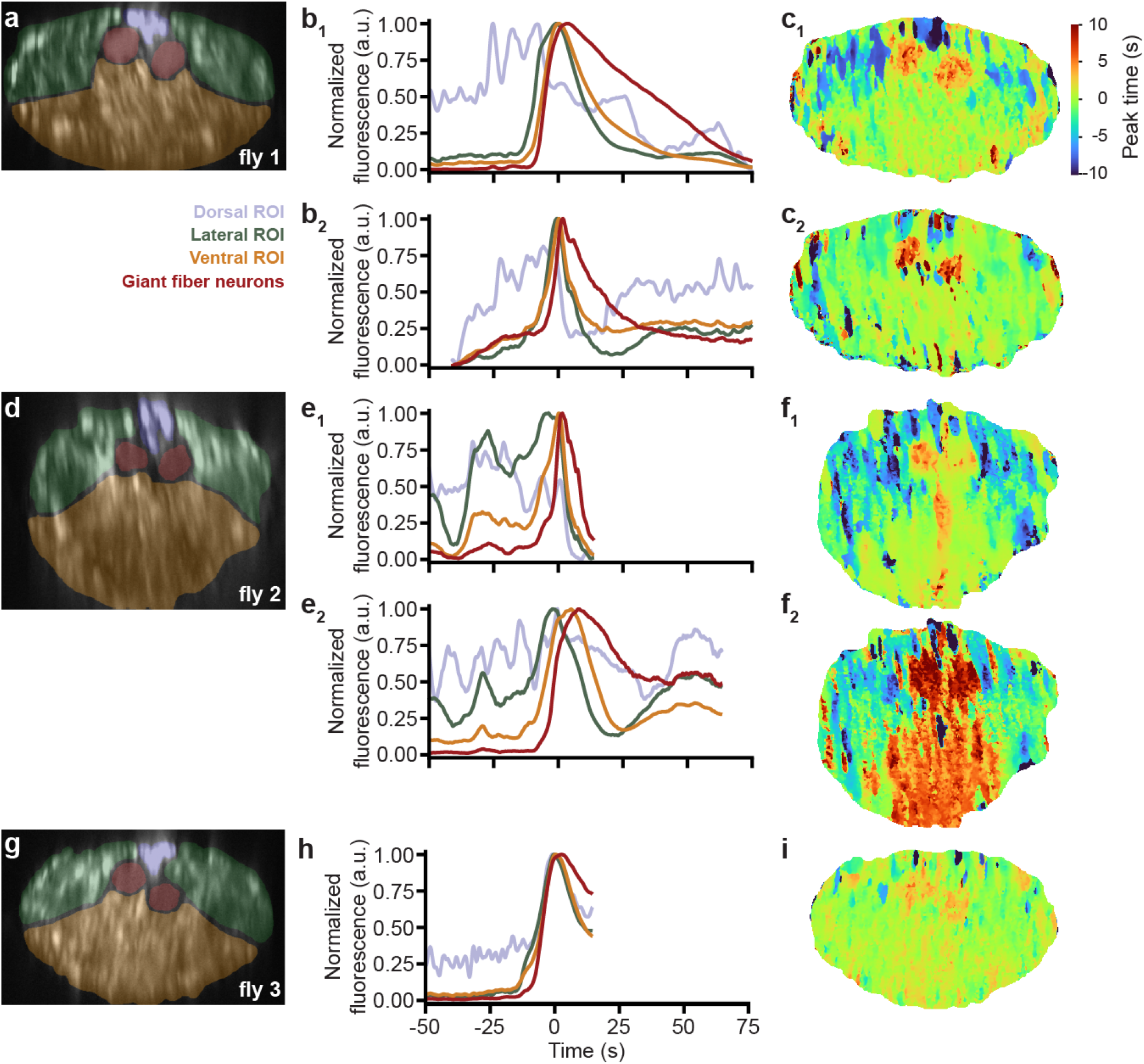
Waves of neural activity observed across three animals following ingestion of high-concentration caffeine. Thoracic cervical connectives from three animals. **(A,D,G)** ROIs overlaid on top of standard-deviation time-projected images. **(B,E,H)** Neural activity over time for each ROI (color-coded) normalized to the peak fluorescence during the wave of activity. Shown are five waves from three animals. Time is aligned to the peak of the mean fluorescence across all ROIs. **(C,F,I)** Pixel-wise time of peak activity (color-coded) relative to the peak of mean activity across the entire neck connective.

## 7 Supplementary Videos

**Video 1: Interactions among implanted and intact freely behaving animals**. Two implanted animals—identifiable by visible thoracic windows—and one intact animal interact near a morsel of food. Video is real-time. https://www.dropbox.com/s/b5ui7z7uotrnoql/video_1.mov?dl=0

**Video 2: Protocol to prepare animals for long-term neural recordings**. A step-by-step visualization of how a fly is outfitted with an implant and window for long-term two-photon microscope recordings. https://www.dropbox.com/s/tpegdzdu80tno4x/video_2.mov?dl=0

**Video 3: Repeatedly recording VNC anatomy across one month**. Two-photon z-stacks of an animal’s VNC at 1, 14, and 28 days post-implantation (dpi). This animal expressed GFP throughout the nervous system (*GMR57C10-Gal4*). Z-stack images progress from the dorsal to ventral VNC. https://www.dropbox.com/s/efntyidl1gnx5aw/video_3.mov?dl=0

**Video 4: Repeatedly recording VNC neural activity across ten days**. Two-photon imaging of an animal’s VNC at 1, 5, and 10 days post-implantation (dpi). This animal expressed a genetically-encoded calcium indicator, GCaMP6s, throughout the nervous system (*Act88F:Rpr; GMR57C10-Gal4; UAS-GCaMP6s*). Neural data are averaged across three cumulatively acquired two-photon microscope images. Activity are related to foreleg-dependent grooming. https://www.dropbox.com/s/4z3bztl88rwm9sl/video_4.mov?dl=0

**Video 5: Optogenetically elicited backward walking in intact, sham implanted, and implanted animals**. Representative videos of three flies driven to walk backward through optogenetic activation of Moonwalker Descending Neurons. Columns are experimental dates (1, 14, and 28 dpi). Rows are experimental groups (Intact, Sham implanted, and Implanted). A light appears on the bottom-left of each arena, indicating times of orange light illumination and CsChrimson activation. Trajectories are shown for forward walking (cyan) and backward walking (purple). https://www.dropbox.com/s/05x5cekrut9gec5/video_5.mov?dl=0

**Video 6: Repeatedly recording the anatomy of proprioceptive inputs to the VNC for 15 days before and after forelimb amputation**. Two-photon z-stacks of two animals’ VNCs at 1, 7, and 15 days-post-implantation (dpi). These animals expressed GFP in limb chordotonal organs (*iav-Gal4*). Z-stack images progress from the dorsal to ventral VNC. Top row shows data from an animal with an intact leg. Bottom row shows an animal whose front left leg was amputated at 2dpi. https://www.dropbox.com/s/lmxsl323qhprots/video_6.mov?dl=0

**Video 7: Repeatedly recording thoracic cervical connective neural activity before, during, right after, and long after feeding with a sucrose solution**. Two-photon imaging of a cross-section of the thoracic cervical connective including neurons descending from and ascending to the brain. Columns are data acquired before (left), during (middle-left), right after (middle-right), and 25 minutes (right) after feeding with a sucrose solution. Rows are behavioral videography (top), Δ*F/F* (middle) and motion-corrected raw (bottom) two-photon calcium imaging data. This animal expressed GCaMP6s and tdTomato, throughout the nervous system. https://www.dropbox.com/s/7zzb2n4570m6ris/video_7.mov?dl=0

**Video 8: Repeatedly recording thoracic cervical connective neural activity before, during, right after, and long after feeding with a low-concentration caffeine and sucrose solution**. Two-photon imaging of a cross-section of the thoracic cervical connective including neurons descending from and ascending to the brain. Columns are data acquired before (left), during (middle-left), right after (middle-right), and 25 minutes (right) after feeding with a low-concentration caffeine and sucrose solution. Rows are behavioral videography (top), Δ*F/F* (middle) and motion-corrected raw (bottom) two-photon calcium imaging data. This animal expressed GCaMP6s and tdTomato, throughout the nervous system. https://www.dropbox.com/s/rn8cas5lxtnyzxs/video_8.mov?dl=0

**Video 9: Repeatedly recording thoracic cervical connective neural activity before, during, right after, and long after feeding with a high-concentration caffeine and sucrose solution**. Two-photon imaging of a cross-section of the thoracic cervical connective including neurons descending from and ascending to the brain. Columns are data acquired before (left), during (middle-left), right after (middle-right), and more than 25 minutes (right) after feeding with a high-concentration caffeine and sucrose solution. Rows are behavioral videography (top), Δ*F/F* (middle) and motion-corrected raw (bottom) two-photon calcium imaging data. This animal expressed GCaMP6s and tdTomato, throughout the nervous system. https://www.dropbox.com/s/28qcd329mhykeu6/video_9.mov?dl=0

**Video 10: Neural activity waves following high-concentration caffeine ingestion**. Two-photon imaging of a cross-section of the thoracic cervical connective including neurons descending from and ascending to the brain. Columns are different occurrences of neural activity waves observed across three animals more than 25 minutes after feeding with a high-concentration caffeine and sucrose solution. Rows are behavioral videography (top), Δ*F/F* (middle) and motion-corrected raw (bottom) two-photon calcium imaging data. These animals expressed GCaMP6s and tdTomato, throughout the nervous system. https://www.dropbox.com/s/84abk0emwsm4klz/video_10.mov?dl=0

## 8 Code and data availability

Code are available at: https://github.com/NeLy-EPFL/Long-Term-Imaging-VNC-Drosophila

Data are available at: https://dataverse.harvard.edu/dataverse/long_term_imaging_vnc_drosophila

## 10 Acknowledgments

We thank Adam Friedberg for help in testing earlier versions of the implant and the manipulation arm. We thank Alain Herzog for photos and videos of implanted animals.

## 12 Author Contributions

L.H. - Conceptualization, Methodology, Software, Validation, Formal analysis, Investigation, Data acquisition, Data curation, Writing - Original Draft Preparation, Writing - Review & Editing, Visualization.

M.K. - Methodology, Software, Validation, Formal analysis, Writing - Original Draft Preparation, Writing - Review & Editing, Visualization.

J.B. - Methodology, Software, Formal analysis, Data curation, Writing - Review & Editing, Visualization.

V.L.R. - Methodology, Software, Formal analysis, Data curation, Writing - Review & Editing, Visualization.

C.-L. C. - Data acquisition, Writing - Review & Editing

S.G. - Methodology, Software, Data curation, Writing - Review & Editing.

F.A. - Methodology, Software, Writing - Review & Editing.

M.S.S. - Conceptualization, Methodology, Resources, Writing – Original Draft Preparation, Writing - Review & Editing, Supervision, Project Administration, Funding Acquisition.

P.R. - Conceptualization, Methodology, Resources, Writing – Original Draft Preparation, Writing - Review & Editing, Supervision, Project Administration, Funding Acquisition.

## References

[1] Denk, W. et al. Anatomical and functional imaging of neurons using 2-photon laser scanning microscopy. Journal of neuroscience methods 54, 151–162 (1994).

[2] Trachtenberg, J. T. et al. Long-term in vivo imaging of experience-dependent synaptic plasticity in adult cortex. Nature 420, 788–794 (2002).

[3] Kim, T. H. et al. Long-term optical access to an estimated one million neurons in the live mouse cortex. Cell reports 17, 3385–3394 (2016).

[4] Andermann, M. L. et al. Chronic cellular imaging of entire cortical columns in awake mice using microprisms. Neuron 80, 900–913 (2013).

[5] Goldey, G. J. et al. Removable cranial windows for long-term imaging in awake mice. Nature protocols 9, 2515 (2014).

[6] Huang, C. et al. Long-term optical brain imaging in live adult fruit flies. Nature Communications 9, 872 (2018).

[7] Seelig, J. D. & Jayaraman, V. Neural dynamics for landmark orientation and angular path integration. Nature 521, 186–191 (2015).

[8] Pick, S. & Strauss, R. Goal-driven behavioral adaptations in gap-climbing drosophila. Current Biology 15, 1473–1478 (2005).

[9] Asahina, K. Neuromodulation and strategic action choice in drosophila aggression. Annual review of neuroscience 40, 51–75 (2017).

[10] Pavlou, H. J. & Goodwin, S. F. Courtship behavior in drosophila melanogaster: towards a ‘courtship connectome’. Current opinion in neurobiology 23, 76–83 (2013).

[11] Seelig, J. D. et al. Two-photon calcium imaging from head-fixed drosophila during optomotor walking behavior. Nature methods 7, 535–540 (2010).

[12] Maimon, G., Straw, A. D. & Dickinson, M. H. Active flight increases the gain of visual motion processing in drosophila. Nature neuroscience 13, 393–399 (2010).

[13] Grover, D., Katsuki, T. & Greenspan, R. J. Flyception: imaging brain activity in freely walking fruit flies. Nature methods 13, 569–572 (2016).

[14] Valle, A. F., Honnef, R. & Seelig, J. D. Automated long-term two-photon imaging in head-fixed walking drosophila. bioRxiv (2021).

[15] Nelson, N. A., Wang, X., Cook, D., Carey, E. M. & Nimmerjahn, A. Imaging spinal cord activity in behaving animals. Experimental neurology 320, 112974 (2019).

[16] Wu, W. et al. Long-term in vivo imaging of mouse spinal cord through an optically cleared intervertebral window. bioRxiv (2021).

[17] Chen, C.-L. et al. Imaging neural activity in the ventral nerve cord of behaving adult drosophila. Nature communications 9, 1–10 (2018).

[18] Tsubouchi, A. et al. Topological and modality-specific representation of somatosensory information in the fly brain. Science 358, 615–623 (2017).

[19] Tuthill, J. C. & Azim, E. Proprioception. Current Biology 28, R194–R203 (2018).

[20] Bidaye, S. S., Bockemühl, T. & Büschges, A. Six-legged walking in insects: how cpgs, peripheral feedback, and descending signals generate coordinated and adaptive motor rhythms. Journal of neurophysiology 119, 459–475 (2018).

[21] Günel, S. et al. Deepfly3d, a deep learning-based approach for 3d limb and appendage tracking in tethered, adult drosophila. Elife 8, e48571 (2019).

[22] Bidaye, S. S., Machacek, C., Wu, Y. & Dickson, B. J. Neuronal control of drosophila walking direction. Science 344, 97–101 (2014).

[23] Klapoetke, N. C. et al. Independent optical excitation of distinct neural populations. Nature methods 11, 338–346 (2014).

[24] Sen, R. et al. Moonwalker descending neurons mediate visually evoked retreat in drosophila. Current Biology 27, 766–771 (2017).

[25] Bavelier, D., Levi, D. M., Li, R. W., Dan, Y. & Hensch, T. K. Removing brakes on adult brain plasticity: from molecular to behavioral interventions. Journal of Neuroscience 30, 14964–14971 (2010).

[26] Sugie, A., Marchetti, G. & Tavosanis, G. Structural aspects of plasticity in the nervous system of Drosophila. Neural Development 13, 14 (2018).

[27] Ayaz, D. et al. Axonal injury and regeneration in the adult brain of drosophila. Journal of Neuroscience 28, 6010–6021 (2008).

[28] Hollis, E. R. Axon guidance molecules and neural circuit remodeling after spinal cord injury. Neurotherapeutics 13, 360–369 (2016).

[29] Hunt, R. F., Scheff, S. W. & Smith, B. N. Synaptic reorganization of inhibitory hilar interneuron circuitry after traumatic brain injury in mice. Journal of Neuroscience 31, 6880–6890 (2011).

[30] Murphy, T. H. & Corbett, D. Plasticity during stroke recovery: from synapse to behaviour. Nature Reviews Neuroscience 10, 861–872 (2009).

[31] Isakov, A. et al. Recovery of locomotion after injury in Drosophila melanogaster depends on proprioception. Journal of Experimental Biology 219, 1760–1771 (2016).

[32] Mamiya, A., Gurung, P. & Tuthill, J. C. Neural coding of leg proprioception in drosophila. Neuron 100, 636–650 (2018).

[33] Root, C. M. et al. A presynaptic gain control mechanism fine-tunes olfactory behavior. Neuron 59, 311–321 (2008).

[34] French, A. S., Geissmann, Q., Beckwith, E. J. & Gilestro, G. F. Sensory processing during sleep in drosophila melanogaster. Nature 1–4 (2021).

[35] Hindmarsh Sten, T., Li, R., Otopalik, A. & Ruta, V. Sexual arousal gates visual processing during drosophila courtship. Nature 595, 549–553 (2021).

[36] Hoopfer, E. D., Jung, Y., Inagaki, H. K., Rubin, G. M. & Anderson, D. J. P1 interneurons promote a persistent internal state that enhances inter-male aggression in drosophila. Elife 4, e11346 (2015).

[37] Gibson, W. T. et al. Behavioral responses to a repetitive visual threat stimulus express a persistent state of defensive arousal in drosophila. Current Biology 25, 1401–1415 (2015).

[38] Shaw, P. J., Cirelli, C., Greenspan, R. J. & Tononi, G. Correlates of sleep and waking in drosophila melanogaster. Science 287, 1834–1837 (2000).

[39] Wu, M. N. et al. The effects of caffeine on sleep in drosophila require pka activity, but not the adenosine receptor. Journal of Neuroscience 29, 11029–11037 (2009).

[40] Lin, F. J. et al. Effect of taurine and caffeine on sleep-wake activity in Drosophila melanogaster. Nature and Science of Sleep 2, 221–231 (2010).

[41] Namiki, S., Dickinson, M. H., Wong, A. M., Korff, W. & Card, G. M. The functional organization of descending sensory-motor pathways in Drosophila. eLife 7, e34272 (2018).

[42] Cande, J. et al. Optogenetic dissection of descending behavioral control in drosophila. Elife 7, e34275 (2018).

[43] Tsubouchi, A. et al. Topological and modality-specific representation of somatosensory information in the fly brain. Science 358, 615–623 (2017).

[44] Ribeiro, C. & Dickson, B. J. Sex Peptide Receptor and Neuronal TOR/S6K Signaling Modulate Nutrient Balancing in Drosophila. Current Biology 20, 1000–1005 (2010).

[45] Wyman, R. J., Thomas, J. B., Salkoff, L. & King, D. G. The drosophila giant fiber system. In Neural mechanisms of startle behavior, 133–161 (Springer, 1984).

[46] Feany, M. B. & Bender, W. W. A drosophila model of parkinson’s disease. Nature 404, 394–398 (2000).

[47] Sinha, S. et al. High-speed laser microsurgery of alert fruit flies for fluorescence imaging of neural activity. Proceedings of the National Academy of Sciences 110, 18374LP–18379 (2013).

[48] Savall, J., Ho, E. T. W., Huang, C., Maxey, J. R. & Schnitzer, M. J. Dexterous robotic manipulation of alert adult drosophila for high-content experimentation. Nature methods 12, 657–660 (2015).

[49] Jang, Y.-H., Chae, H.-S. & Kim, Y.-J. Female-specific myoinhibitory peptide neurons regulate mating receptivity in Drosophila melanogaster. Nature Communications 8, 1630 (2017).

[50] Duffy, D. C., McDonald, J. C., Schueller, O. J. & Whitesides, G. M. Rapid prototyping of microfluidic systems in poly(dimethylsiloxane). Analytical Chemistry 70, 4974–4984 (1998).

[51] Johansson, A., Calleja, M., Rasmussen, P. A. & Boisen, A. SU-8 cantilever sensor system with integrated readout. Sensors and Actuators, A: Physical 123–124, 111–115 (2005).

[52] Soft lithography for micro-and nanoscale patterning. Nature Protocols 5, 491–502 (2010).

[53] Weibel, D. B., DiLuzio, W. R. & Whitesides, G. M. Microfabrication meets microbiology. Nature Reviews Microbiology 5, 209–218 (2007).

[54] Laermer, F., Schilp, A., Funk, K. & Offenberg, M. Bosch deep silicon etching: Improving uniformity and etch rate for advanced MEMS applications. Proceedings of the IEEE Micro Electro Mechanical Systems (MEMS) 211–216 (1999).

[55] Satoshi, K., Hong-Bo, S., Tomokazu, T. & Kenji, T. Finer features for functional microdevices. Nature 412, 697–698 (2001).

[56] Liu, Y. et al. Deformation Behavior of Foam Laser Targets Fabricated by Two-Photon Polymerization. Nanomaterials (Basel, Switzerland) 8 (2018).

[57] Sridhar, V. H., Roche, D. G. & Gingins, S. Tracktor: image-based automated tracking of animal movement and behaviour. Methods in Ecology and Evolution 10, 815–820 (2019).

[58] Paszke, A. et al. Pytorch: An imperative style, high-performance deep learning library. In Wallach, H. et al. (eds.) Advances in Neural Information Processing Systems 32, 8024–8035 (Curran Associates, Inc., 2019).

[59] Nair, V. & Hinton, G. E. Rectified linear units improve restricted boltzmann machines. In International Conference on Machine Learning (ICML), 807–814 (2010).

[60] Srivastava, N., Hinton, G., Krizhevsky, A., Sutskever, I. & Salakhutdinov, R. Dropout: a simple way to prevent neural networks from overfitting. The Journal of Machine Learning Research 15, 1929–1958 (2014).

[61] Kingma, D. P. & Ba, J. Adam: A method for stochastic optimization. The International Conference on Learning Representations (ICLR) (2015).

[62] Schindelin, J. et al. Fiji: an open-source platform for biological-image analysis. Nature methods 9, 676–682 (2012).

[63] Distinctive image features from scale-invariant keypoints. International Journal of Computer Vision 60, 91–110 (2004).

[64] Lecoq, J. et al. Removing independent noise in systems neuroscience data using DeepInterpolation. Nature Methods (2021).

